# Dysregulation of the secretory pathway connects Alzheimer’s disease genetics to aggregate formation

**DOI:** 10.1101/2020.08.10.243634

**Authors:** Chih-Chung Kuo, Austin WT Chiang, Hratch M. Baghdassarian, Nathan E. Lewis

## Abstract

A hallmark of amyloid disorders, such as Alzheimer’s disease, is aggregation of secreted proteins. However, it is largely unclear how the hundreds of secretory pathway proteins contribute to amyloid formation. We developed a systems biology framework that integrates expression data with protein-protein interaction networks to successfully estimate a tissue’s fitness for producing specific secreted proteins. Using this framework, we analyzed the fitness of the secretory pathway of various brain regions and cell types for synthesizing the Alzheimer’s disease-associated amyloid-precursor protein (APP). While none of the key amyloidogenic pathway components were differentially expressed in AD brain, we found the deposition of Aβ is associated with repressed expression of the secretory pathway components proximal to APP. Concurrently, we detected systemic up-regulation of the secretory pathway components proximal to β- and γ-secretases in AD brains. Our analyses suggest that perturbations from 3 high confidence AD risk genes cascade through the secretory machinery support network for APP and into the endocytosis pathway. Thus, we present a model where amyloidogenesis is associated with dysregulation of dozens of secretory pathway components supporting APP, which could yield novel therapeutic targets for the treatment of AD.

## Introduction

No mammalian cell exists alone. Indeed, each cell dedicates >1/3 of its protein-coding genes to interact directly with other cells and its environment (Uhlén et al., 2015, 2019), using hormones and receptors for communication, enzymes and other proteins to modify their extracellular matrix, transporters for exchanging metabolites, etc. The mammalian secretory pathway is tasked with the synthesis, post-translational modification (PTM), quality control, and trafficking of these secreted proteins (secPs) (Novick et al., 1981; Reynaud and Simpson, 2002). SecPs account for >25% of the total proteome mass(Hukelmann et al., 2016; Tan et al., 2017), and are among the most tissue-specific genes in the human genome (Uhlén et al., 2015). The precision and efficiency of the mammalian secretory pathway result from the concerted effort of hundreds of secretory machinery components (secMs) including chaperones, enzymes, transporters, glycosyltransferases, metabolites and lipids within the secretory pathway (Feizi et al., 2013, 2017; Gutierrez et al., 2018; Lund et al., 2017). Since many secPs relay signals between cells or modify a cell’s microenvironment, each cell must carefully regulate the synthesis and localization of each secP.

Perturbations to the secretory pathway result in misfolded proteins, which induce ER stress and apoptosis. In amyloid diseases, the misfolded proteins can aggregate into toxic amyloid fibrils, which ultimately lead to cell death. Aβ deposition, a major pathological hallmark of Alzheimer’s disease (AD), stems from the perturbed processing of the transmembrane amyloid precursor protein (APP). In Aβ aggregation, alternative proteolytic cleavage of amyloid precursor peptide by β-(rather than α-secretase) releases the secreted form of APP, the aggregation-prone Aβ1-42.(De Strooper et al., 1998; Lammich et al., 1999; Vassar et al., 1999) Furthermore, additional PTMs in the secretory pathway may affect APP cleavage (Thinakaran and Koo, 2008; Wang et al., 2017), phosphorylation (Lee et al., 2003), glycosylation (Joshi and Wang, 2015; McFarlane et al., 1999, 2000; Schedin-Weiss et al., 2014) and trafficking (Jiang et al., 2014; Wang et al., 2017). However, protein aggregation could stem from the perturbation of diverse processes, but no systematic exploration of all processes supporting proper APP processing has been done (Knowles et al., 2014). Several large-scale GWAS also identified more than 45 AD risk loci (Dourlen et al., 2019; Jansen et al., 2019; Kunkle et al., 2019), although for many loci, it remains unclear how they induce Aβ deposition. Furthermore, a large part of AD heritability remains unknown (Dourlen et al., 2019; Ridge et al., 2016). The genetic landscape of late-onset Alzheimer’s (LOAD) is highly heterogeneous, with multiple complex molecular interactions contributing to the disease phenotype. Therefore, the discovery of concerted expression changes in LOAD, such as the remodeling of immune-specific modules requires systems approaches on large datasets (Zhang et al., 2013).

To unravel the molecular changes leading to Aβ deposition, we focused on the roles of the secretory pathway in amyloidogenesis. The secretory pathway is responsible for the processing, quality control and trafficking of key components of the amyloidogenic pathway (Greenfield et al., 1999; Hartmann et al., 1997), such as APP and the secretases, so we investigated if there is a systemic dysregulation of the secMs supporting their production and processing. To do this, we first developed a network-based approach that leverages protein-protein interaction (PPI) and mRNA and protein abundance data to quantify a cell or tissues’ “secretory machinery support”. This measures the fitness of a tissue or cell for properly secreting a specific secP based on the expression of its supporting secMs. Next, we investigated if there are disruptions in the secretory machinery support for key players of the amyloidogenic pathway (i.e., APP and the secretases), leading to increased Aβ deposition in LOAD, based on data from several large-scale clinical bulk- and single-cell RNA-Seq datasets (Mathys et al., 2019; Wang et al., 2018). We found significant dysregulation of the secretory pathway proximal to APP and the secretases, and this dysregulation is a major determinant of Aβ deposition. We further demonstrated that the concerted expression changes in the secretory support modules for the APP, BACE1, and PSEN1 can be linked to known AD risk genes and their regulation targets. In terms of subcellular localization, the core perturbed network enriches for known hotspots for Aβ production such as ER, cytosol and endosomes. Moreover, we found that the AD risk loci activate endocytosis via the core support network, and we identified a candidate TF binding motif that is conserved in the promoter regions of the interaction network genes. Together, our analyses suggest mechanisms underlying impaired protein secretion, which could propose novel therapeutic targets for the treatment of AD. It also proposes mechanisms by which AD genetics imbalance the secretory pathway, thus resulting in Aβ deposition, cell death, and cognitive impairment.

## Results

### Secreted proteins and secretory machinery show similar tissue-specific expression

The secretory pathway synthesizes and transports a variety of secreted proteins, each with different requirements for their synthesis and secretion (e.g., different physicochemical properties and post-translational modifications). With the human secretome being one of the most tissue-specific subsets of the human proteome (Ramsköld et al., 2009; Uhlén et al., 2015), we hypothesized each tissue expresses just the secMs needed to synthesize and process secPs from the tissue. Supporting this, we observed that clustering tissues by secP gene expression grouped tissues similarly as when clustering by secM gene expression (Figure 1, p-value = 0.0145; Figure S1). Thus, the secMs are not merely housekeeping proteins always expressed to support any proteins being secreted; rather, they express in a tissue-specific fashion to meet the demands of different tissues (Feizi et al., 2017). However, the question remains if the pairings between secMs expressed in each tissue represent those needed to specifically support the secPs they secrete.

**Figure 1.**
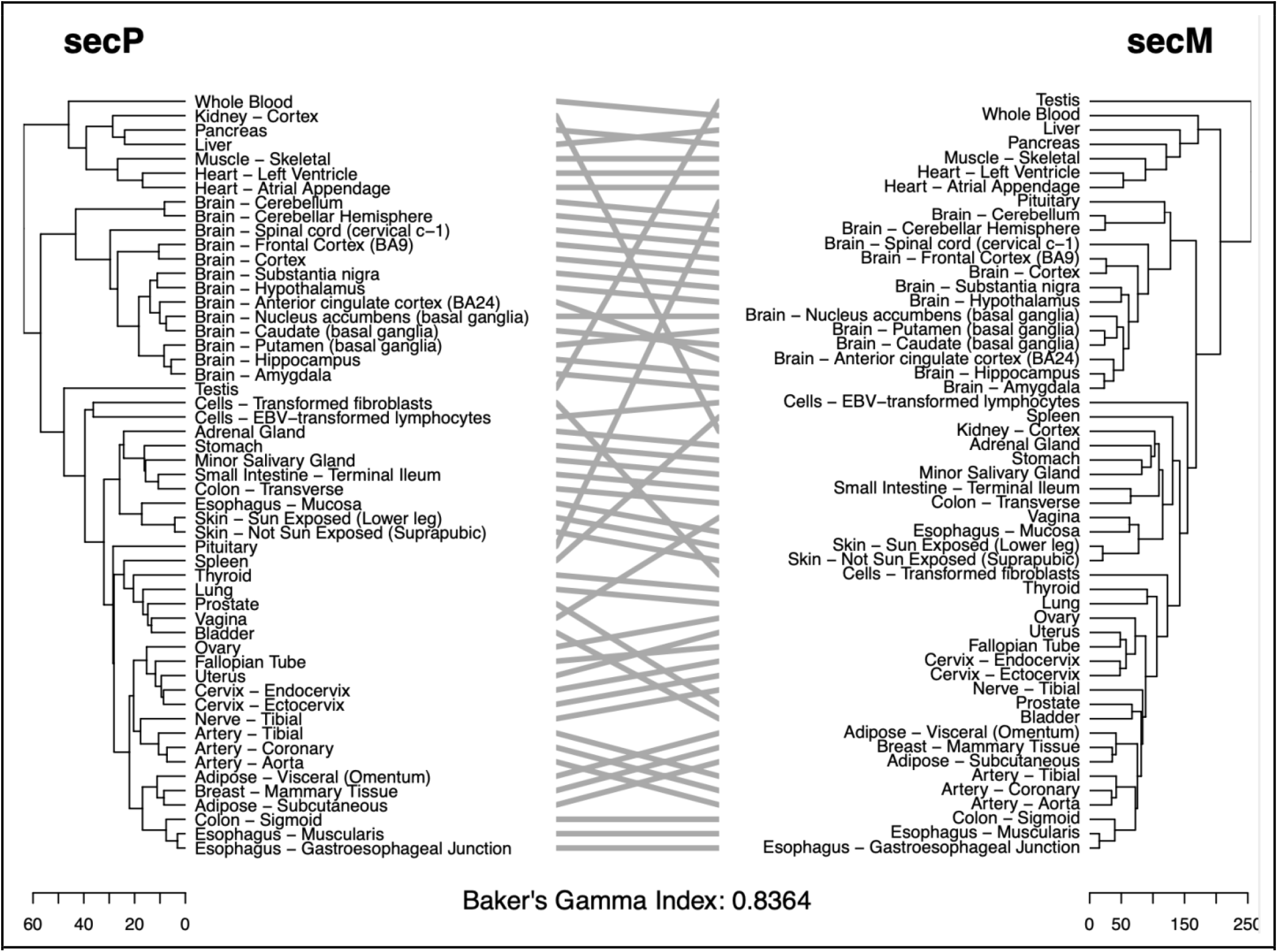
The secMs and the secPs show coordinated expression profiles across different human tissues. The tissue similarity structures from the perspective of the secretome and the secretory pathway expression were represented by two hierarchical clustering dendrograms, which were then compared with a tanglegram (Galili, 2015). Gene expression of the secretory pathway and its clients show a high level of coordination across human tissues, with precision significantly better than expected by sampling genes from each tissue (bootstrapping p-value 0.013 and 0.045 for data from GTEx (GTEx Consortium, 2015) and Human Protein Atlas (Uhlén et al., 2015) respectively).

### Tissue specific expression of secMs predict expression of their client-secreted proteins

To further dissect the pattern of secP-secM co-regulation seen across tissues, we incorporated two sources of information: protein-protein interactions (PPIs) and secM gene expression. We harness PPIs to identify the secMs relevant to each secP, since PPIs are one of the major modalities through which machinery proteins in the secretory pathway assist protein secretion (Anelli and Sitia, 2008; Bonifacino and Glick, 2004; Ikawa et al., 1997; Pearl and Prodromou, 2006). Further, secMs responsible for secP post-translational modifications are well-captured by PPIs between the secPs and the secMs (Figure S2). To focus on spatially proximal interactions, we filtered the PPIs for interactions between secMs and other secretory pathway-resident proteins for each secP (see Methods), resulting in a “secM support network” consisting of 3658 genes. By overlaying secM gene expression on this network, one can quantify the secM support for secretion of each secP. To systematically quantify the fitness of the secM support network for producing each secP in a tissue, a machinery support score is calculated for each secP by a random walk algorithm that integrates secM gene expression levels proximal to each secP in its PPI network. More specifically, for each secP, we added the protein to the secM support network, and centered the network on the secP. We then performed a random walk on the secM support network starting from the secP. We adapted the transition probabilities of the random walk to incorporate gene expression of the secMs so that propagation is constrained by not only PPI network topologies but also the expression of the secM components, allowing one to contextualize cell- and disease-specific interactomes (methods and supplementary note; Figure 2a). The algorithm assigns a component score to each protein in the network, representing its availability to the secP of interest. The “secretory machinery support score” (i.e., the average component scores the secretory pathway components receive from the random walk) then quantifies the overall secretory pathway support for the given secP. Using this approach, we found the secretory machinery support score for each secP increases in tissues wherein the secP is more highly expressed (Figure 2b, Figure S3). Further, the machinery support score considerably improves the prediction of secP protein abundance from mRNA expression across tissues (Figure S4 and supplementary note). Thus, by accounting for the mRNA/protein expression of PPIs surrounding each secP, the machinery support score quantifies a tissue’s relative fitness for synthesizing and secreting the secP of interest.

**Figure 2.**
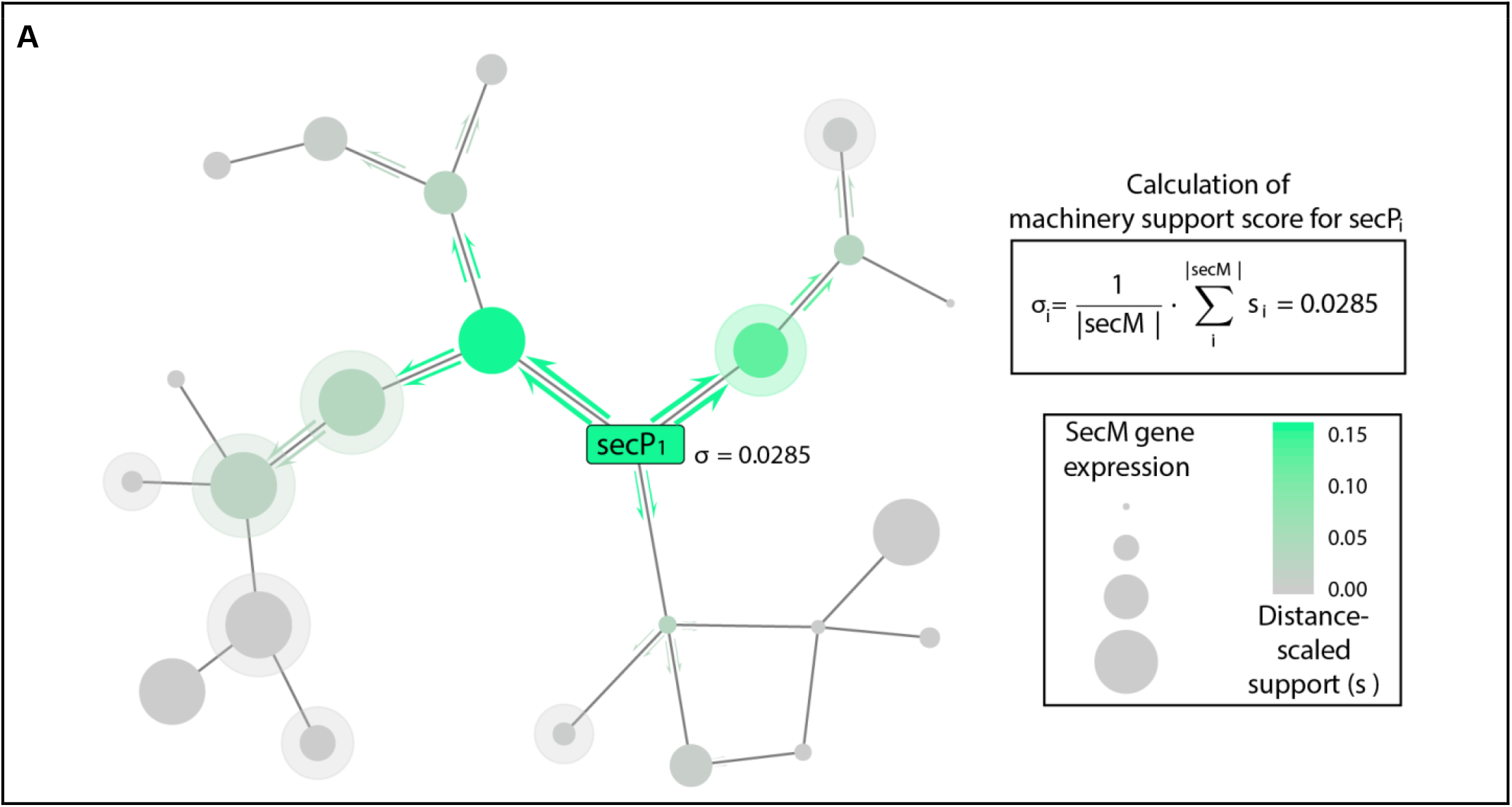
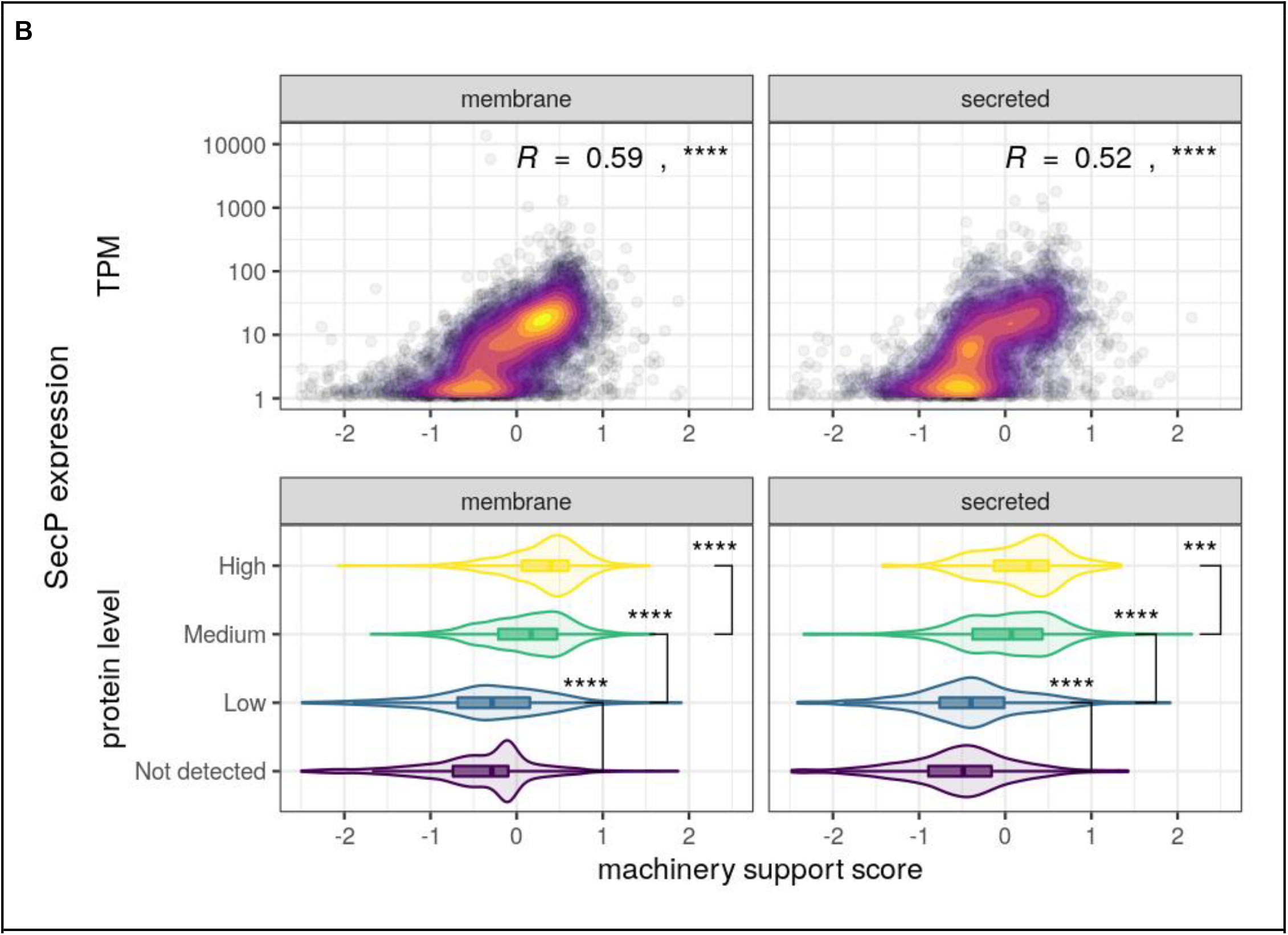
Expression data can quantify a tissue or cell’s fitness for synthesizing a secreted or membrane protein. **(A)** In our systems biology approach, the mRNA or protein abundance is overlaid on a PPI network surrounding a secreted protein (secP). The secP synthesis fitness is quantified by summing the secM expression, scaled by distance from the secP (computed by a random walk), yielding a quantitative “machinery support score”. The calculation of the support score also provides a sub-network of proteins contributing to secP synthesis. **(B)** We quantified the machinery support score for every secreted protein in all tissues in the Human Protein Atlas, and found a clear correlation between the secP expression and the relative machinery support score. This correlation was seen for both mRNA (top, spearman correlation coefficient, see methods) and protein (bottom, t-test) abundance. Thus, our machinery support score allows one to quantify how fit a cell or tissue is for properly expressing and processing a secreted or membrane protein.

### Aβ deposition in Alzheimer’s disease is characterized by perturbed secretory support of amyloid precursor protein

The co-regulation of the secP and their cognate secMs results from millions of years of evolution. Thus, the question arises whether perturbations to such co-regulation could underlie the molecular pathology in AD. Specifically, Aβ deposition is a major hallmark of AD-pathology. The precursor to Aβ, APP, is moderately to highly expressed (high transcript levels and moderate protein levels) in the cerebral cortex. While APP overexpression from APP duplication can cause early-onset (familial) AD (Bushman et al., 2015; Rovelet-Lecrux et al., 2006), sporadic (non-familial) AD does not show differential transcript abundance for APP between AD and non-AD individuals despite the increase in Aβ plaques (Matsui et al., 2007). However, APP does undergo post-transcriptional processing, with pathogenic Aβ being released from APP following sequential cleavage by β- and γ-secretases, while the α-secretase promotes the correct processing of APP.

To test the relevance of secretory pathway expression to AD, we analyzed RNA-Seq data from 4 brain regions in 298 AD and age-matched control subjects (from the Mount Sinai Brain Bank (Wang et al., 2018)) and single-cell RNA-Seq from the prefrontal cortex (Brodmann area 10) of 48 individuals (Mathys et al., 2019) (Figure 3). APP was not differentially expressed in AD brains at both the single-cell (Mathys et al., 2019) and the tissue level (Wang et al., 2018). This is not surprising since the amyloidogenic pathway giving rise to neurotoxic Aβ takes place post-translationally (De Strooper et al., 2010). Additionally, expression of neither BACE1 nor PSEN1 correlated with plaque abundance in affected brain regions (Figure 3). However, each gene had significant changes in machinery support scores correlating with severity (Figure 3b). Specifically, the supporting machinery score for APP decreased in affected brain regions, showing suppressed scores in cells with amyloid deposition (p<0.0066). The largest effect was in cell types that are major producers of Aβ, including neurons (Greenfield et al., 1999; Hartmann et al., 1997; Laird et al., 2005), reactive astrocytes (Frost and Li, 2017; Liddelow and Barres, 2017; Phatnani and Maniatis, 2015; Sofroniew and Vinters, 2010) (Figure 3a, Figure S5), and in brain regions affected early in the onset of AD (Figure 3b, Figure S6). However, BACE1 and PSEN1, which aid in amyloidogenesis, showed an opposite trend, with affected cells increasing machinery support for these secretases (Figure 3b; p-values for Brodmann area 36 (parahippocampal gyrus, BM36) and Brodmann area 44 (inferior frontal gyrus, BM44): p <0.0038 and p<0.014 for β-secretase; p<0.13 and p<0.083 for γ-secretase).

**Figure 3.**
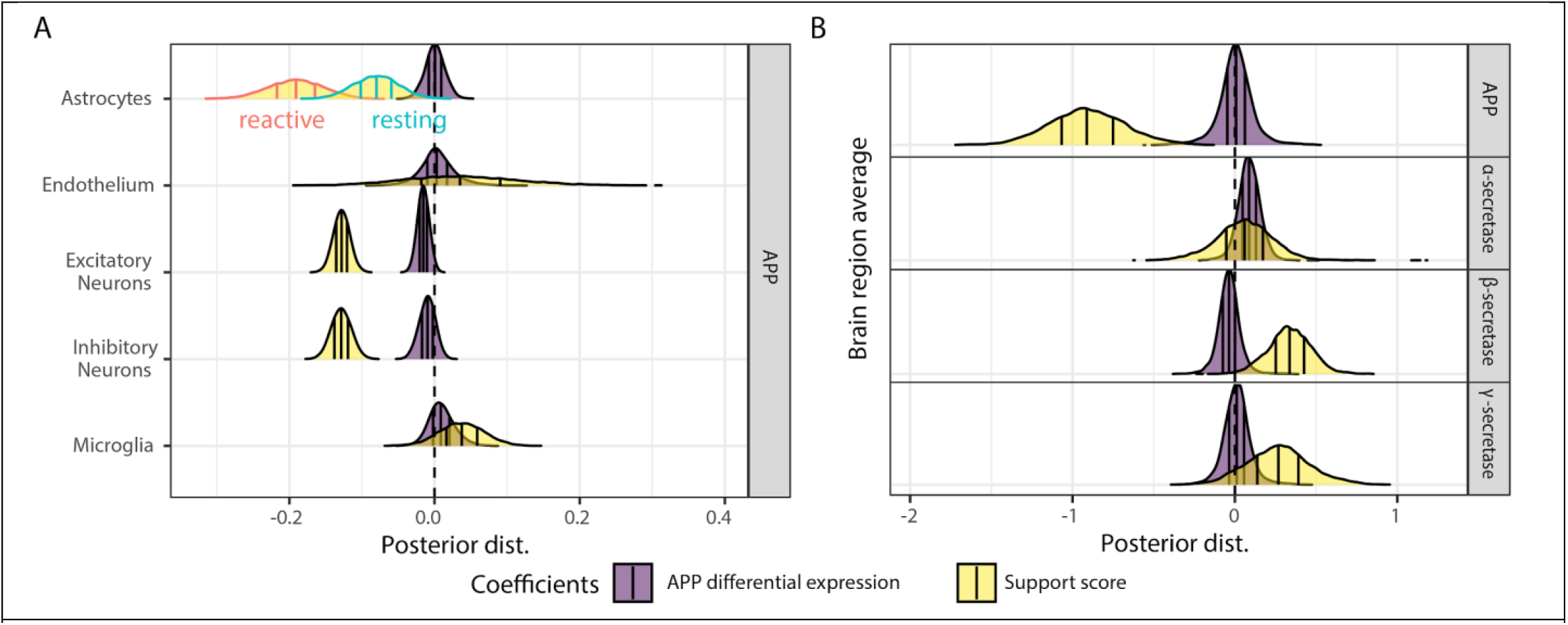
AD-relevant genes show perturbed secretory machinery support scores. **(A)** Overall, APP expression does not correlate with plaque densities across individuals in single-cell RNA-Seq. However, support scores show a negative correlation with plaque density, suggesting the secMs supporting proteostasis of APP are suppressed in AD. **(B)** Similar trends were seen in all brain regions surveyed (BM10, BM22, BM36, BM44, Figure S5). On average, no correlation is found between plaque abundance and gene expression of APP, BACE1 or PSEN1. However, secM support score for APP shows a negative correlation while the support scores for BACE1 and PSEN1 positively correlate with amyloid formation, suggesting AD pathogenesis involves dysregulation of the secretory pathway.

### Secretory pathway support of APP is most strongly suppressed proximal to APP

The secretory machinery support score for APP is significantly decreased in AD. However, it is unclear if the decline in APP support score is due to a general suppression of many secMs throughout the secretory pathway or a local repression in which only the secMs most proximal to APP are down-regulated. To test this, we defined a core support network for APP based on network proximity and gene expression. As the abundance and proximity to APP vary across the secMs in the network, we broke down the APP support score into the individual component scores for each protein in the secM support network (Table S1). This quantifies each secM’s corresponding contribution to the secretory support of APP. When the secMs are rank-ordered by their individual component scores, their contribution to the APP support score follows a pattern of exponential decay (Figure S7A), suggesting that the support score is mostly determined by a smaller number of proteins with high proximity to APP (Figure 4A). While the entire APP support network is not differentially expressed between AD and healthy brains, we wonder if this is the case with secMs that are major contributors to the support score. When we overlaid the differential expression across brain regions and cell types and considered progressively smaller subsets of the APP support network consisting of proteins with the highest component scores, we saw the strongest repression at around 20-30 secMs, suggesting that the proteins nearest to APP are the most suppressed (Figures 4A, S7B, S7C).

**Figure 4.**
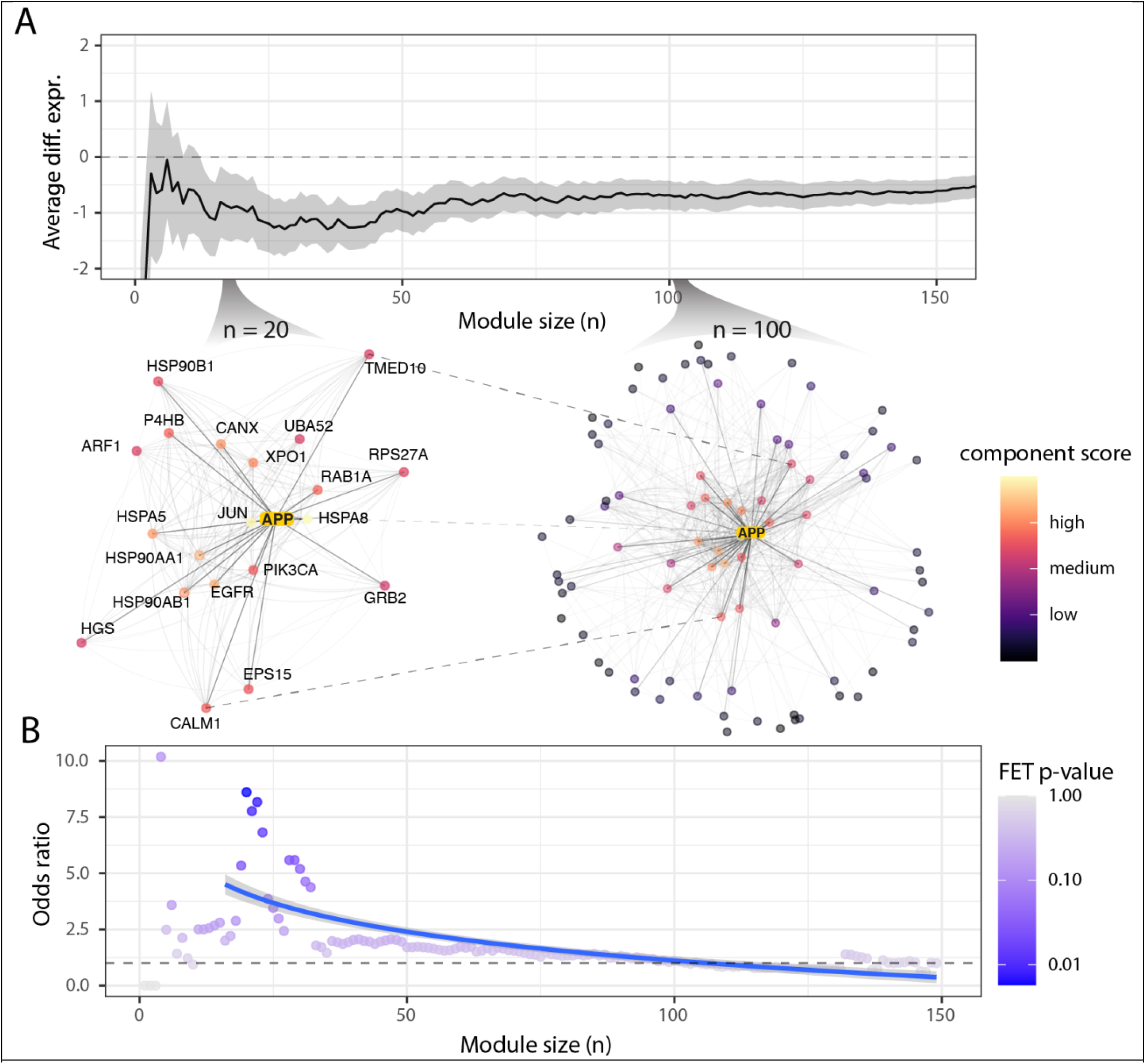
The APP secM proteostasis network is not enriched for AD risk genes, but is enriched for AD risk gene regulatory targets. The networks of secM proteins supporting APP production at two support component score cutoffs, representing the top *n* = 20 / *n* = 100 proteins contributing the most to the APP’s support score. The top 20 proteins with the highest component scores are labeled. **(A)** Starting from n=1 where the secM with the highest component score was considered, we incrementally included secMs less proximal to APP. The sizes of the subnetworks are indicated by the x-axis. At each iteration, we calculated the average differential expression (y-axis) between AD and healthy subjects for each subnetwork. The strongest repression of the AD support network occurs at around n=15∼30. **(B)** We also calculated the degree to which the subnetwork enriched for regulatory targets of known AD risk genes above the genome-wide significance threshold (y-axis). The regulatory targets of AD risk genes are generally depleted among the general non-APP-specific secMs (Figure S9 for full trend across all 3685 subnetworks), but targets of AD risk genes enrich strongly among the core secMs closest to APP.

### Changes in APP-supporting PPIs are regulated by AD risk loci

Large GWAS screens found AD risk genes impacting many pathways (Kunkle et al., 2019; Lambert et al., 2013). Although the secretory pathway is tasked with the synthesis and processing of APP, it is not generally implicated in LOAD pathogenesis, since secretory pathway genes are not enriched among LOAD risk genes from large-scale GWAS studies (Kunkle et al., 2019). However, our results show transcriptional perturbations of machinery support for key amyloidogenic genes; thus, there may be a concerted regulatory change for modules supporting APP production and proteostasis in the secretory pathway. Thus, we tested if regulatory AD risk loci (i.e., transcription factors) regulate the secMs interacting with APP. While the entire APP support network does not enrich for the AD risk genes nor their regulatory targets, we wondered if AD risk loci selectively target the secMs more proximal to APP in the PPI network. As we assessed enrichment for AD risk gene targets in subnetworks that were progressively closer to APP (i.e., more focused on the support network interacting most directly with APP), we found the secMs supporting APP were increasingly enriched for targets of AD risk regulatory genes (Figures 4B, S8, S9). The results are reproducible across multiple GWAS significance thresholds (Figure S9). Since the enrichment of AD risk gene targets steadily increased in statistical significance with increasing proximity to APP in the PPI network, we focused on the top 20 proteins interacting the most closely with APP from the support network. This resulted in an APP support subnetwork where the enrichment significance for AD risk gene targets peaks. This cutoff also coincides with the subnetwork with the strongest suppression of secM expression (Figures 4A, S7B, S7C).

### Core support network overlaps significantly with genomic loci with differential histone acetylation in AD brain

Epigenetic alterations have been linked to neurodegeneration in human AD brains and AD mouse models (Lardenoije et al., 2015; Liu et al., 2018). Thus, we analyzed data from three epigenome-wide association studies to investigate if the core support network is overrepresented for hotspots of aberrant epigenomic reprogramming in AD. We found that proteins proximal to APP on the support network show significant enrichment for Aβ related epigenetic changes measured in H3K9ac profiles from 669 aged human prefrontal cortices (Klein et al., 2019) (Figure 5, top row). Additionally, the enrichment around the core network is higher for H3K9ac peaks annotated as being in enhancer domains than those in promoter domains. In another epigenome-wide association study comparing aging- and AD-related histone acetylation changes (Nativio et al., 2020a), we found the H3K122ac, H3K27ac and H3K9ac peaks that differ significantly were disproportionately located near the core support network (Figure S10, top row). Interestingly, while AD and aged brains often share similar epigenetic signatures (Nativio et al., 2018), enrichment of AD-related peaks (Figure S10, bottom row) is stronger in the core support network than that of aging-related peaks (Figure S10, middle row). Thus, we observe considerable epigenetic changes in human AD brain focused around the APP supporting network.

**Figure 5.**
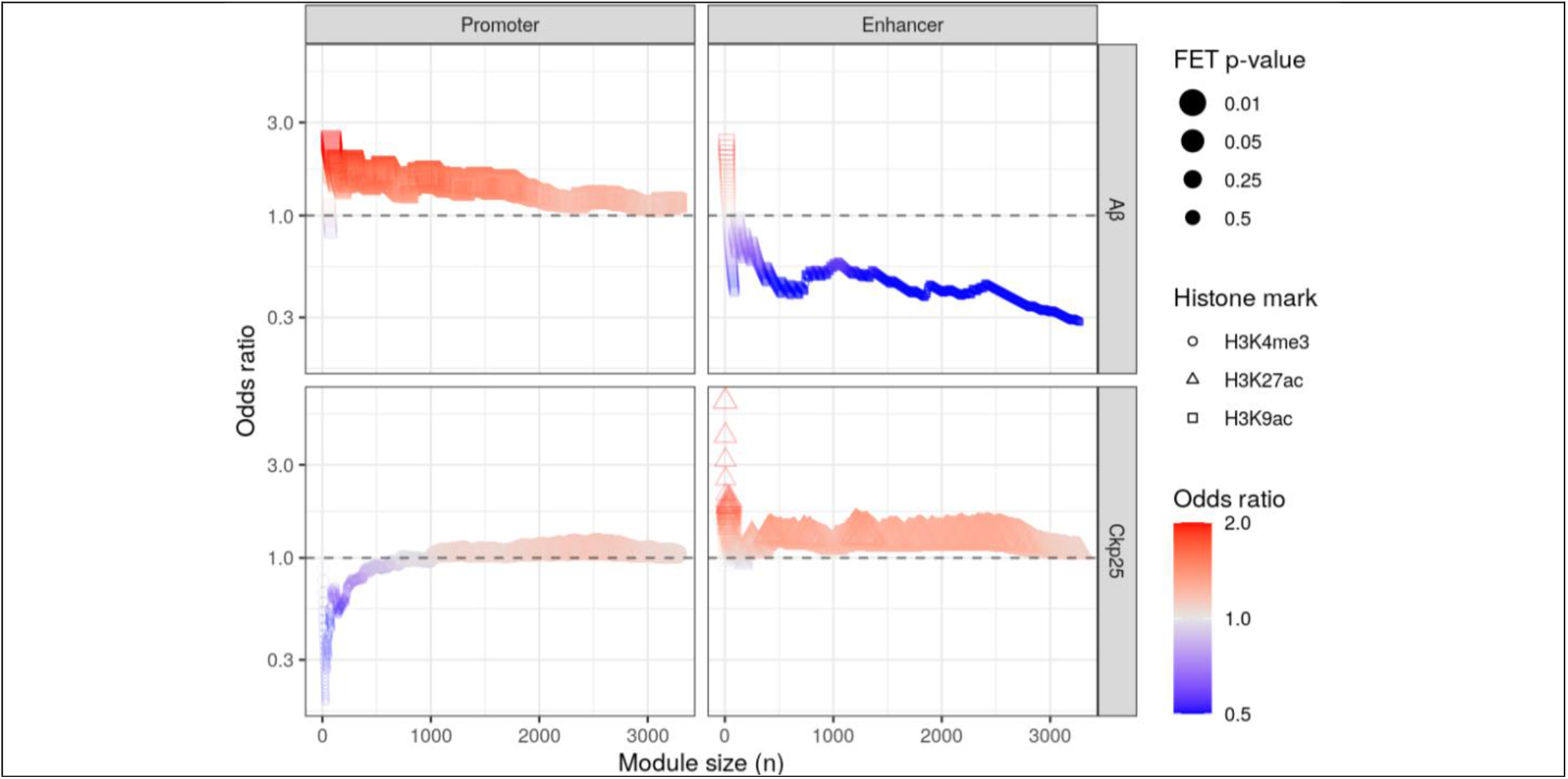
The APP core support network is enriched for genes whose enhancer regions contain AD-specific histone marks. Following the notations from Figure 4, in each subplot the y-axis (odds ratio) shows the degree to which the core support network overlaps with genomic loci with differential histone modifications is indicated evaluated at modules of various sizes (x-axis), with a smaller n indicating a smaller subnetwork with only the proteins most proximal to APP. The subplots are arranged based on the region in which the differentially-enriched peaks are detected (columns), and the conditions compared (top row, Aβ-associated changes; bottom row, CK-p25 mice with increased Aβ levels vs. controls).

The enriched epigenetic changes were further captured in a mouse model of AD. Specifically, histone methylation and acetylation marks were profiled in CK-p25 mice with increased Aβ levels and controls (Gjoneska et al., 2015). While the core network is depleted for significantly altered H3K4me3 peaks in CK-p25 mice, AD-associated H3K27ac alterations are significantly enriched among proteins proximal to APP (Figure 5, bottom row). This is in line with our previous observation in which the core support network is a hotspot for AD-related acetylation marks, especially around the enhancer domains.

### AD risk loci activate endocytosis via the core support network

We analyzed the content of the core support network, and found it is enriched for genes in the amyloidogenic pathway. Specifically, we saw interactions were concentrated in the ER, endosomes, and the cytosol (Figure S11, FDR p<1.1e-3), consistent with the localization of amyloidogenesis. For example, the endosome hosts intracellular Aβ production with its β-secretase, and is enlarged in autopsies from AD (Cataldo et al., 2000) and stem cell models (Israel et al., 2012). To further unravel the link between endocytosis and the core support network, we analyzed the patterns of the core network differential expression between AD and controls across multiple cohort studies using gene regulatory networks obtained from ENCODE (ENCODE Project Consortium, 2012) and Ingenuity Pathway Analysis (IPA) (Krämer et al., 2014). Complementing our previous observation that a significant portion of the core support network is endosome-resident, we saw significant up-regulation of genes associated with endocytosis (p-value = 8.95e-14), mediated by the core supporting machinery (Figures 6A, S12) across various brain regions.

**Figure 6.**
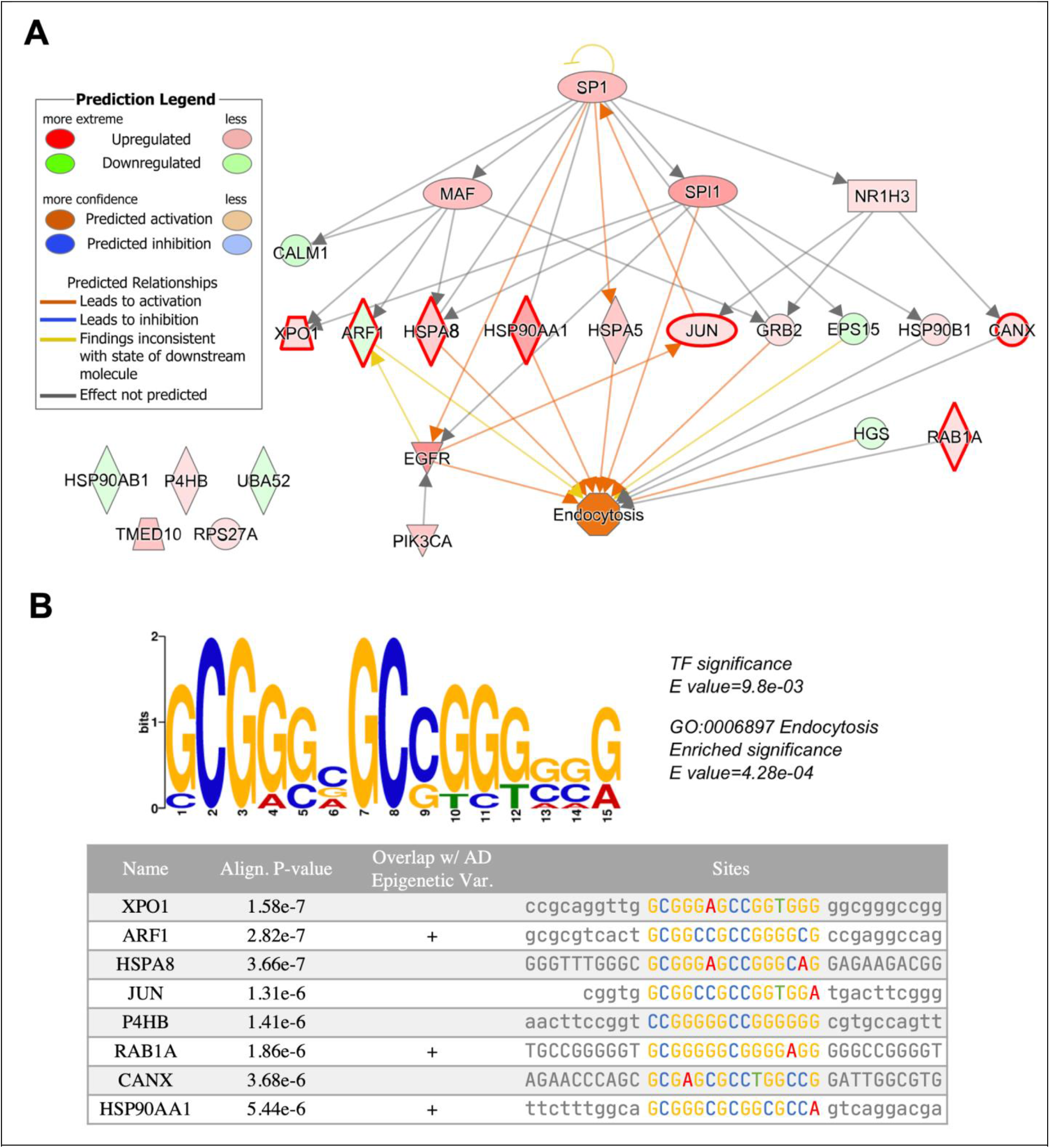
The regulatory relationships surrounding the core support network. (A) Regulatory structures of the 20 proteins from the core support network were constructed using IPA. Gene expression profiles from the 4 brain regions (BM10, BM22, BM36, BM44) (Wang et al., 2018), the Mayo Study (Allen et al., 2016) and the ROSMAP study (Religious Order Study and Memory and Aging Project) (De Jager et al., 2018) were averaged and overlaid on the core support network. The endocytosis pathway is strongly activated in AD brains via the core support network, which in turn is regulated by 3 genes from AD risk loci--NR1H3, MAF and SPI1. The proteins harboring binding sites for the TF motif (shown in panel B) were outlined in red. (B) The TF motif (top panel) aligns significantly to 8 genes from the entire support network (bottom panel), 7 of which belong to the core support network. Align. P-value, statistical significance of the motif alignment to the promoter region of each gene; overlap w/AD epigenetic Var., whether the motif binding site completely overlaps with significant epigenetic alterations in AD.

We further analyzed the APP core support network and identified transcription factors (TFs) that are most strongly associated with the perturbed secM module. The APP core support network coincides significantly with the regulatory targets of 3 genes from AD risk loci: NR1H3, MAF and SPI1 (Dourlen et al., 2019; Jansen et al., 2019; Kunkle et al., 2019; Lambert et al., 2013). To investigate the extent to which these AD risk genes perturb transcription of the supporting machinery in AD, we analyzed the differentially expressed genes between AD and controls across multiple cohort studies using gene regulatory networks obtained from ENCODE (ENCODE Project Consortium, 2012) and Ingenuity pathway analysis (IPA) (Krämer et al., 2014).

In addition to predicting upstream transcriptional regulators associated with amyloidogenesis from gene signatures or differentially expressed genes, we searched for conserved TF binding sites among the top-ranked dysregulated genes of the core-network in Alzheimer’s Disease. Specifically, we conducted *de novo* TF binding site motif discovery for the 20 genes in the core support network (Figure 6B; see Methods). We identified one significant TF motif (E value= 9.8e-3) that is present in 7 of the genes in the core support network (Figure S13), which is associated with the SP1, SP2, and SP3 transcription factors. SP1 is important in AD (Citron et al., 2015; Santpere et al., 2006) and is predicted to bind the 3 genes from AD risk loci (NR1H3, MAF, and SPI1) with high confidence via three elite enhancers (Fishilevich et al., 2017) GH11J047390 (GH score=2.2), GH16J079764 (GH score=2.4), and GH11J047247 (GH score=2.1) respectively (Table S5). Furthermore, several motif binding sites across the core support network significantly overlap with loci with major epigenetic alterations in AD (Figure 6B) (Klein et al., 2019; Nativio et al., 2018, 2020b). For example, the motif binding site at the promoter region of ARF1 completely overlaps with a histone acetylation mark H3K122ac that is significantly altered in AD but not aged subjects (chr1:228259654-228280877, Wilcoxon Rank Sum P-value 0.004). Another locus of significantly altered H3K122ac peaks in AD individuals (chr2:65353363-65359754, Wilcoxon Rank Sum P-value 0.01) overlaps with the motif binding site at RAB1A. The motif binding site at HSP90AA1 overlaps with significantly altered peaks for H3K122ac and H3K9ac, which are repressed and upregulated respectively in AD (chr14:102543639-102575586, Wilcoxon Rank Sum P-value 0.05 and chr14:102543639-102575586, Wilcoxon Rank Sum P-value 0.05). Thus, a perturbation to multiple TFs could disrupt the APP core support network, wherein both genetic risk genes (NR1H3, MAF, and SPI1) and global regulators can contribute to the dysregulation of the core network.

Further analysis suggests the AD risk genes and the core support network genes are further co-regulated with an activation of endocytosis in the AD pathogenesis. We further characterized the functional association of the conserved TF motif by scanning all promoters of genes in the genome for the motif (see Methods for details). The conserved TF motif was significantly enriched in the promoter regions of known endocytic pathway genes (p-value=4.28e-4) and several other pathways relevant to AD pathogenesis (Table S6, Figure S11). Together, these results suggest a concerted change in endosomal activities and dysregulated pathways between normal and AD brains that arises due to the differential expression of the core supporting machinery surrounding APP.

## Discussion

In the past four decades, the Alzheimer’s research community has made huge strides towards elucidating molecular processes contributing to amyloid deposition. However, despite involving aggregation of secreted proteins, it remained unclear to what extent the core processes of protein secretion and proteostasis are involved. Here we developed a systems biology approach that analyzed the interactions between key amyloidogenic components and the secretory pathway. This approach predicted the propensity for amyloid deposition at the single-cell level. To gain systems-level insights into LOAD, we used the framework to identify a subset of the secretory pathway components on which concerted suppression and several regulatory elements of AD converged. We further demonstrated that an increase in endocytic activities in LOAD can be attributed to key AD risk genes via the core support network.

LOAD is a complex disease. The identification of three rare mutations in APP, PSEN1 and PSEN2 that occur in early-onset familial AD and the discovery of APOE constitute our primary knowledge of the genetic landscape of LOAD. More recently, high-throughput technologies such as GWAS and whole exome sequencing have identified more than 45 genetic risk loci of LOAD. However, the additional risk loci exert only very small risk effects (Bertram et al., 2010), and the link between genetic risk variants and amyloid deposition remains incompletely understood. Despite the highly heterogeneous expression of APP and other key amyloidogenic components across AD and healthy subjects, we demonstrated concerted down-regulation of secretory machinery proximal to APP in AD patients. This highlights the secretory pathway as a determinant of amyloid deposition, which had not been a major focus of AD research. Incidentally, the proteostasis network, with which the secretory pathway shares a significant overlap, has been an increasingly popular target of protein aggregation and aging studies. The human chaperone network, a major component of the proteostasis network, undergoes continual remodeling during an organism’s lifespan (Brehme et al., 2014; Hipp et al., 2019; Walther et al., 2017). However, in the aging AD brain, the directions of regulation of these chaperones are rather uncoordinated across different chaperone families and even within the same family (Brehme et al., 2014). Our observations of concerted repression of key proximal secretory pathway components show that improper expression of the secretory pathway, of which the chaperone network is a subset, is associated with the deposition of amyloid.

Even though the secretory pathway is in charge of post-translational processing and targeting of APP up until its cleavage by the secretases, its implications in AD have been insufficiently researched primarily due to the lack of AD risk genes in the pathway (MacArthur et al., 2017). Our results showed that genes contributing the most to the APP support network in the secretory pathway are significantly enriched for targets of AD risk genes and AD related epigenetic changes, suggesting a mechanistic link between genetic and epigenetic variants of AD and secretory pathway dysregulation, complementing previous systems-level approaches to understanding LOAD with a focus on immune-and microglia-specific modules (Zhang et al., 2013). To further unravel the link, we examined the core support network consisting of secretory pathway components most proximal to APP. We noticed 3 AD risk genes that are also transcription factors showing regulatory evidence over the core support network according to both curated and *de novo* pathway analyses. More importantly, we demonstrated a regulatory cascade originating from the 3 AD risk genes, mediated by the core support network and into the endocytosis pathway. The endosome, where the β-secretase is localized to and where its acidic pH is optimal for enzymatic cleavage, is a major site of intracellular Aβ production. Our findings thus offer a mechanistic view of amyloidogenesis involving the secretory pathway and the endosomes. This is in line with observations in embryonic cortical neurons that showed increased Aβ levels as a result of increased endocytic pathway activities and reuptake in APP in aged cells (Burrinha et al., 2019). We also observed significant enrichment of endosomal-localized proteins in the core support network for APP, further lending credence to the involvement of the secretory pathway in activating the endocytic pathway.

The dominant model of AD pathogenesis, the amyloid hypothesis (Hardy and Higgins, 1992; Hardy and Selkoe, 2002; Selkoe and Hardy, 2016), posits that AD pathogenesis and the rest of the disease process such as tau tangle formation (Hardy et al., 1998; Lewis et al., 2001) result from the accumulation of Aβ via the imbalance between Aβ production and clearance. We examined the capacity of the secretory pathway in the context of Aβ production and processing, where the secretory support of APP and β- and γ-secretases were analyzed. While the concerted dysregulation of the secretory support for these key amyloidogenic components in AD brains theoretically leads to increased Aβ production, the impact on Aβ clearance warrants further investigation. It is worth noting that detectable Aβ deposition can precede the onset of AD by more than 15 years (Bateman et al., 2012; Jack et al., 2013), which likely coincides with the onset of the decline of the proteostasis network. Our findings highlight the roles of the secretory pathway in amyloidogenesis, which open new possibilities for early diagnosis and treatment research on LOAD. Furthermore, our systems approach can be further applied to other diseases in which the secretory pathway is perturbed, such as perturbed hormone secretion in endocrine disorders, changes in hepatokine secretion nonalcoholic fatty liver disease (Gorden et al., 2015; Meex et al., 2015), and the secretion of diverse proteins in cancer (Robinson et al., 2019).

## Methods

### Calculation of secretory pathway support scores

#### Random walk on interactome

To contextualize the secretion of a given secP, we used network propagation to quantify the influence of gene expression across neighboring genes. Let *G*(*V, E*) denote an undirected interactome with vertex set *V* containing *n* proteins and an edge set *E* the *m* interactions between them. Let *w*_*ij*_ be the edge weight (*w*_*ij*_ *=* 1 if *G* is undirected) between edges *i* and *j* and *A* be the adjacency matrix of *G* where *A*_*ij*_ *=* {*w*_*ij*_ *if* {*v*_*i*_,*v*_*j*_} ∈ *E*; 0 *otherwise*}. Let *x*(*t*)be the location of the walk at time *t*. Note that given the previous walk location at time *t* − 1, we can represent the probability of the walker moving from location *i* to *j* in a single step as:

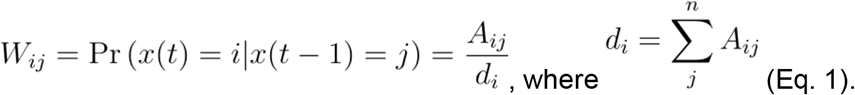

Summing probabilities from all inbound locations we have :

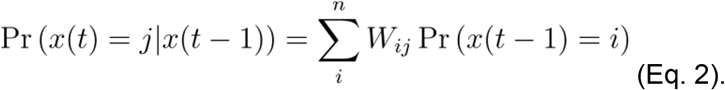

In matrix notation, this is **p**(*t*) = **Wp**(*t* − 1), where W is the transition matrix and each entry *W*_*ij*_ denotes the aforementioned transition probability from *i* to *j*. In random walk with restarts (Page et al., 1998), at each step the walk resets to the origin with probability α, and the last equation becomes **p_RWR_**(t) = (1 − α) **Wp**_**RWR**_(*t* −1) + α**P**(0), where **P**(0) = *e*_*k*_ denotes the initial distribution if the walker starts at *v*_*k*_. The restart parameter α was set to 0.1, as advised by the linear optimal model given the size of the network (Huang et al., 2018).

#### Expression-guided random walk with restarts

The transition matrix can be modified to incorporate gene expression into each step of the propagation. If we let ***t*** = [*t*_*0*_,…,*t*_*n*_]^T^, where *t*_*i*_ is the scaled expression corresponding to node,*v*_*i*,_*0* ≤ *t*_*i*_ ≤ 1, the expression-adjusted transition matrix can thus be given by 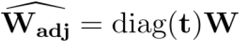. The choice of **t** can either be protein levels where available or mRNA abundance. We show the validity of using mRNA abundance as input in the supplementary notes. We can normalize the adjusted transition matrix by adding in self-loop: 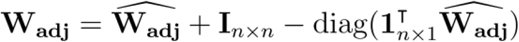,and the update rule, which we termed expression-guided random walk with restarts (eRWR), now becomes:

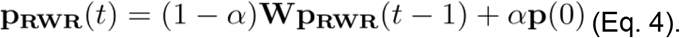

### Calculation of support scores and component scores

We performed eRWR on each secP of interest for 20 iterations (Supplementary Notes), and the final vector of probabilities represent the support component score for each gene on the network *G*(*V, E*). The support score (σ) is the average of the support component scores of the secretory pathway proteins,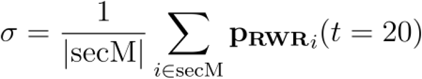

#### Context filtering of the secretory pathway support network

We used the composite consensus human interactome PCNet v1.3 (Huang et al., 2018) (NDEX UUID: 4de852d9-9908-11e9-bcaf-0ac135e8bacf) to be the static, context-agnostic network (*G*_0_(*V*_0_,*E*_0_)). For each secP, we create a subnetwork containing the secP and other essential secretory pathway genes by filtering the network for human secretory pathway components (Feizi et al., 2017; Gutierrez et al., 2018) and secretory pathway-resident proteins (Thul et al., 2017) to constrain the network spatially, resulting in a vertex-induced subgraph *G*(*V =*{*V*_0_ ∩ (*secP* ∪ *secM* ∪ *secResident*)},{*uv*|*uv* ∈ *E*_0_ *and u,v* ∈ *V*}).

#### Transcriptomic and proteomic data processing and support scores calculation for normal human tissues

To calculate support scores for the normal human secretome, we used two datasets, the Human Protein Atlas (HPA) (Uhlén et al., 2015), and the deep proteome and transcriptome abundance atlas (deep proteome) (Wang et al., 2019) for tissue-specific transcriptomes from healthy human donors in which matching proteomic data were also available. For data from the Human Protein Atlas, we downloaded the transcriptomic data--“RNA HPA tissue gene data” and performed log- and sigmoid-transformation on the transcript abundance (TPM) data, resulting in transformed gene expression profiles in the (0,1)range. For the HPA dataset, the support score was calculated based on the tissue-median of the transformed gene expression profiles for each secP. We retained the semi-quantitative nature of the immunohistochemistry protein abundance reporting, and we calculated the support scores summary statistics for proteins belonging to each of the staining levels--”High”, “Medium”, “Low” & “Not detected” separately.

With the fully quantitative proteomic data from the deep proteome (Wang et al., 2019), We calculated the support scores based on the protein iBAQ values. The iBAQ abundance values were transformed in a similar fashion as the transcript abundance from the HPA dataset. Namely, they were log- and then sigmoid-transformed into the (0,1)range, before being median-summarized by tissue and subsequently used in the calculation of the support scores.

#### Transcriptomic and proteomic data processing and support scores calculation for AD and healthy brains

To calculate support scores for key amyloidogenic pathway components in AD and healthy brains, we used two major transcriptomic datasets from the ROSMAP project (Religious Orders Study and Memory Aging Project)--single-cell (Bennett et al., 2018; Mathys et al., 2019) (Synapse ID syn18485175) and bulk RNA-seq (Wang et al., 2018) (Synapse ID syn3159438) data from individuals respectively with varying degrees of Alzheimer’s disease pathology. The single-cell transcriptomic dataset covers 80660 cells from the prefrontal cortex of 48 individuals. While annotations for major cell types were given, we further classified astrocytes into reactive and non-reactive astrocytes based on GFAP expression (Liddelow and Barres, 2017). The bulk RNA-seq covers 4 brain regions (Brodmann areas 10, 22, 36, 44) of 364 individuals.

To transform the count data into appropriate expression inputs to the eRWR algorithm, count data from healthy tissue-specific transcriptomes and the AD single-cell RNA-seq data were first log-transformed to compress the extreme values. The values were then z-score standardized and passed through a logistic function, where the final transformed values have a range (0,1). For the AD bulk-RNA seq data, since the counts were already normalized and transformed, they were z-score standardized and transformed by a logistic function without first being log-transformed.

### Statistical analysis

#### Relationship between support scores of secreted proteins and protein expression

To examine the dependencies between support scores and the transcript and protein abundances of the human secretome, we calculated the support score for each secreted protein in the human secretome (Uhlén et al., 2019) across various human tissues. We first calculated the spearman correlation coefficients between the tissue-median support scores and the transcript and protein abundances across all secreted proteins. To assess the statistical significance of the spearman correlation coefficients, a t-statistic 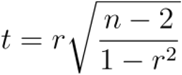 where n and r indicate the number of paired observations and the pearson correlation coefficient respectively was computed. P-values were then calculated by comparing the t-statistic with its null distribution (the t-statistics approximate a t-distribution with n-2 degrees of freedom under the null hypothesis) (Kendall and Stuart, 1977).

To further quantify the statistical significance of transcript abundance and protein level in determining overall protein abundance, a Bayesian hierarchical model was created (model equations shown below) where the abundance of each protein across the 29 tissues is drawn from a linear combination of mRNA levels and the support scores weighted by their respective regression coefficients. We used the rethinking R package(McElreath, 2020) to construct the model and sample the coefficients.

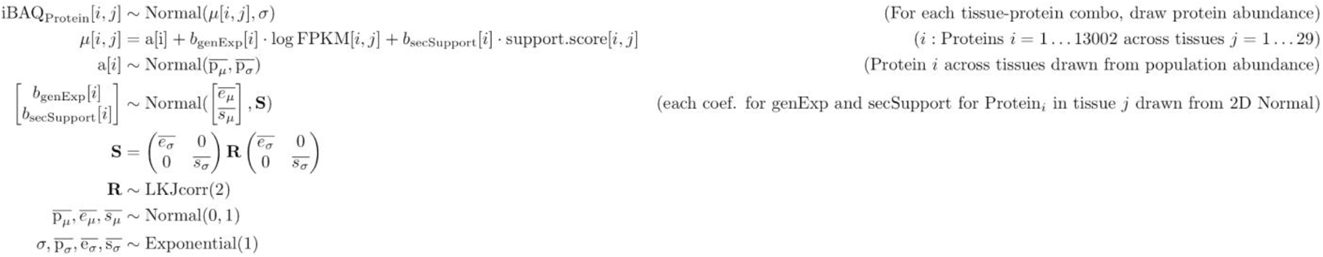

#### Relationship between support scores of key amyloidogenic proteins and amyloid plaque densities

We built a Bayesian hierarchical model (equations shown below) to determine the extent to which the support scores for key amyloidogenic pathway components including APP and the secretases for each cell/ sample in the single-cell (see the supplementary notes for adaptations to the model formula to account for sample covariates) and bulk RNA-seq dataset affects the amount of amyloid plaque measured. We regressed the scaled amyloid plaque densities corresponding to the individual from which the single-cell/ bulk RNA-seq sample was collected against the gene expression and secretory pathway support scores of key amyloidogenic pathway components. To regularize the coefficients of interest, their Bayesian priors are all normally distributed around 0. The coefficients were sampled using the rethinking R package(McElreath, 2020).

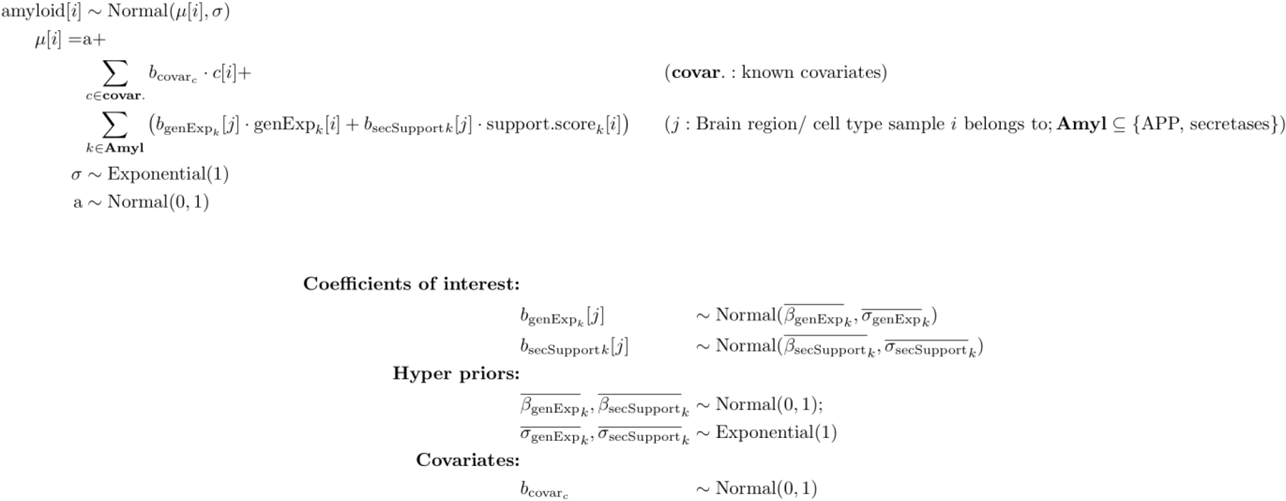

### Characterizing the core support network

#### AD risk genes and enrichment analysis of regulatory components

We obtained 45 genome-wide significant risk loci identified by several AD GWAS studies as summarized previously (Dourlen et al., 2019), resulting in 176 high-confidence AD risk genes. We compiled a separate set of AD risk genes from GWAS summary statistics (Jansen et al., 2019; Kunkle et al., 2019) for loci above the genome-wide suggestive threshold, where MAGMA (de Leeuw et al., 2015) was used to aggregate p-values for SNPs to the gene-level independently for each GWAS dataset. P-values from the two datasets for each gene were then combined using Fisher’s method, resulting in 673 AD suggestive risk genes.

The transcription factors and their targets were obtained from ENCODE (Davis et al., 2018) and ChEA (Lachmann et al., 2010) via the Enrichr portal (Kuleshov et al., 2016). To determine whether the core support network enriches for the regulatory targets of AD risk genes, we first calculated the level of overlap between the core support network and the targets of each transcription factor using Fisher’s exact test, where significantly overlapping transcription factors were defined as those with p-values of less than 0.05. A secondary enrichment was performed to quantify the level to which the significant transcription factors overlap with known AD risk genes. As mentioned earlier, two lists of AD risk genes were used. For the 673 AD suggestive risk genes, a traditional Fisher’s exact test was performed. For the risk genes originating from the 45 risk genome-wide significant risk loci, instead of calculating the direct overlap between the significant transcription factors and the 176 high-confidence risk genes, we mapped the significant transcription factors back to the 45 risk loci on which Fisher’s exact test was performed. This is motivated by the fact that many risk loci contain multiple risk genes that cannot be further pinpointed due to complex linkage disequilibrium patterns, a risk locus is considered hit if at least one of its mapped risk genes appears significantly enriched as a transcription factor. We performed this two-stage enrichment analysis starting from the full static support network towards the core support network by pruning back proteins furthest from APP in each iteration.

#### Enrichment analysis of subcellular compartments

We compiled lists of proteins for all subcellular structures consisting of proteins known to localize to the compartment of interest within the cell (Thul et al., 2017). We ordered the proteins in the full support network by the extent to which they deviate from their stationary support component score to control for network topology while accounting for secretory-resident proteins. To determine the degree to which the proteins from certain subcellular compartments are overrepresented in the core subnetwork, we applied Gene Set Enrichment Analysis (GSEA) (Korotkevich et al., 2016; Subramanian et al., 2005) with the subcellular localization gene-sets and the ranked core support network components as input, eliminating the need for a hard significance cut -off. Subcellular compartments significantly enriched in the core subnetwork are defined as those with an FDR p-value of 0.05 or less.

#### Causal gene network analysis

To robustly define the core supporting subnetwork, we iteratively constructed subnetworks from proteins most proximal to APP and progressively include more distal proteins corresponding to different significance cutoffs. To robustly select the cutoff for the core supporting subnetwork, we performed the two-stage enrichment analysis on all subnetworks as detailed above (see “AD risk genes and enrichment analysis of regulatory components”). Additionally, we calculated the average differential expression between AD and healthy individuals for each subnetwork using fold changes from bulk and single-cell RNA Seq data depending on the source expression from which the subnetwork is calculated. We selected 20 proteins most proximal to APP to include in the final core subnetwork, where the cutoff coincides with the strongest enrichment of regulatory AD risk loci and the suppression of the core subnetwork.

To determine the regulator effects, we performed two network-based analyses. We first ran the upstream regulator analysis using the curated regulator networks from IPA (Krämer et al., 2014). The algorithm took as inputs the core subnetwork and the differential expression fold changes and p-values. Batch-corrected differential gene expression profiles between AD and healthy brains from the Mount Sinai study (Wang et al., 2018), the Mayo Study (Allen et al., 2016) and the ROSMAP study (Religious Order Study and Memory and Aging Project) (De Jager et al., 2018) were obtained from the AMP-AD Knowledge Portal (Synapse ID syn14237651). “Disease & functions” having considerable overlap with the core subnetwork were added, of which endocytosis is the most significant (p-value =2.34E-14).

#### *De novo* TF binding site motifs discovery and known TF binding site identification

We downloaded promoter sequences (version: GRCH38) from UCSC Genome Browser(Kent et al., 2002) for the core subnetwork. The promoter sequences are defined as sequences 1,000 bases upstream of annotated transcription start sites of RefSeq genes with annotated 5’ UTRs. To conduct de novo TF binding site motifs discovery, we first ran motif discovery using the MEME suite(Bailey et al., 2015) with default parameters to identify candidate TF binding site motifs within the promoter sequences by using the entire APP support network serving as background control. Then, the MEME discovered TF binding site motifs were analyzed further for matches to known TF binding sites for mammalian transcription factors in the motif databases, JASPAR Vertebrates (Sandelin et al., 2004), via motif comparison tool, TOMTOM(Gupta et al., 2007). We summarized all the enriched GO terms using ‘Revigo‘(Supek et al., 2011) (Figure S13) on the 81 GoMo identified specific enriched GO terms in the Biological Process (Table S2).

## Data availability

https://github.com/LewisLabUCSD/AD_secretory_pathway

## Acknowledgments

This work was supported by NIGMS (R35 GM119850, NEL), the Ministry of Education (Taiwan), and the Novo Nordisk Foundation (NNF10CC1016517, NNF20SA0066621). Study data were provided by the Rush Alzheimer’s Disease Center, Rush University Medical Center, Chicago. Data Collection was supported through funding by NIA grants P30AG10161, R01AG15819, RR01AG17917, R01AG30146, R01AG36836, U01AG32984, U01AG46152, the Illinois Department of Public Health, and the Translational Genomics Research Institute. ROSMAP data can be requested at https://www.radc.rush.edu.

