## Supplementary Notes and Figures for "Dysregulation of the secretory pathway connects Alzheimer’s disease genetics to aggregate formation"

#### Implementation details for support score calculation

To calculate the expression-guided random walk with restarts (eRWR, see methods) the mRNA or protein expression profile is used as an input along with a protein-protein interaction (PPI) network. Additional tuning parameters include the restart parameter and the number of iterations for which the eRWR is performed. Here, we present the rationale behind the choices of certain input datasets and parameters.

##### Addressing the tissue-specificity of PPI network

The molecular basis of a disease is often tissue or cell-type specific. Thus, one must account for the tissue or cellular context when studying such mechanisms. However, there are very limited disease or even tissue-specific protein interaction data; thus it has been difficult to comprehensively estimate the cell or patient-specific interaction profiles of APP and other amyloidogenic proteins. By integrating transcriptomics and PPIs, the eRWR algorithm accounts for cell type and tissue-specificity by integrating transcriptomic or proteomic data into the PPI. To demonstrate this, we repeated our analysis using cell-type-specific interactomes from the human brain (Mohammadi et al., 2019). We focused on the inhibitory neuron as it is the only cell-type where high-confidence interactions are available for APP. The inhibitory neuron-specific interactions were then paired with expression profiles from neurons (Mathys et al., 2019) to calculate support scores for both inhibitory and excitatory neurons. For the inhibitory neurons profiled, the APP support scores calculated based on the inhibitory neuron-specific network correlated highly with the original support scores derived using the generic interactome (Supplementary Figures N1).

To further demonstrate the robustness of our method, we ran the Bayesian hierarchical model with the updated APP support scores (Supplementary Figures N2), and observed a significant negative association between the support scores with Aβ burden closely mirroring the trend from Figure 3. The concordance between support scores derived from a generic PPI network and those from a cell-type-specific interactome demonstrates our method's ability to contextualize cell- and disease-specific networks.

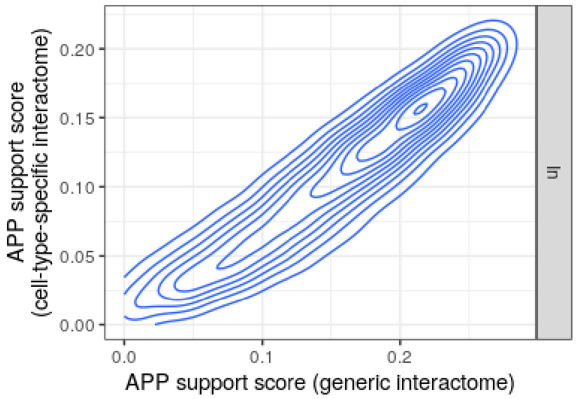


**Figure N1**
The APP support scores from the inhibitory neurons calculated using the generic interactome (x-axis) closely match the scores calculated using the cell-type-specific interactome (y-axis).

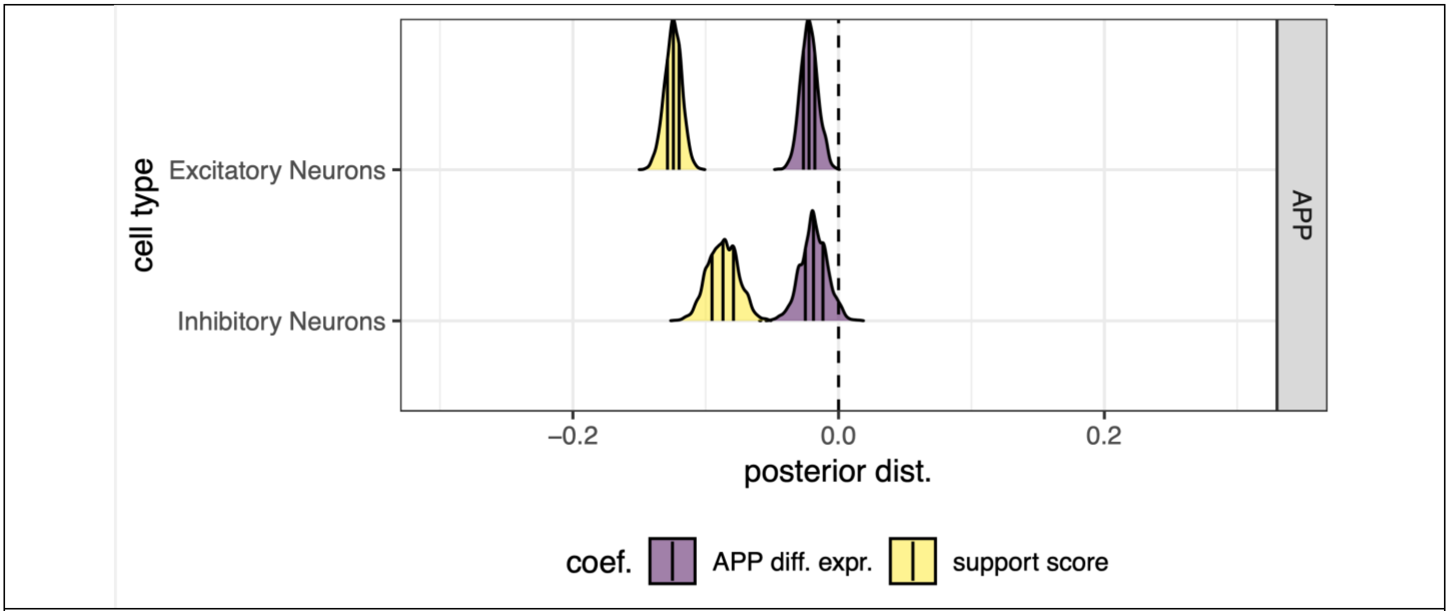


**Figure N2**
The support scores calculated using the neuron-specific interactome show a negative correlation with amyloid deposition, similar to the pattern shown in Fig. 3A.

#### Interactions between partially-folded proteins

As the mammalian secretory pathway is tasked with the synthesis, post-translational modification, quality control and trafficking of secreted proteins (Novick et al., 1981; Reynaud and Simpson, 2002), it is possible that some nascent secreted proteins might not have adopted their mature conformation when interacting with certain secretory pathway machinery. The protein-protein interactions used in this study (Huang et al., 2018) are compiled from measurements using a variety of methods that may include PPIs without necessarily discriminating between intermediate or mature forms of the secPs (Huttlin et al., 2015, 2017; Luck et al., 2020; Oughtred et al., 2019; Rolland et al., 2014; Szklarczyk et al., 2019). It is worth noting, however, in the case of Aβ production, most of the well-known pathogenic processes take place near the end of the secretory pathway (Greenfield et al., 1999; Hartmann et al., 1997). It is expected that secPs near their end of the journey through the secretory pathway will have adopted near steady-state conformation, and these interactions will in turn be captured by the proposed algorithm. However, for other diseases whose pathogenic processes take place earlier along the secretory pathway, the inclusion of transient interactions, potentially at the spatial-temporal resolution, could better help dissect the dynamics of the secP during its early synthesis.

#### Validity of using mRNA expression in place of protein abundance as input to eRWR

In this study, mRNA expression profiles were used for both bulk and single-cell samples of AD and healthy brains since no comparable high-coverage proteomic datasets exist to the best of our knowledge. As mRNA levels are not always the best representative proxies for protein abundance, we demonstrate below that our machinery support score for a given protein in a tissue can be reliably estimated either by mRNA or protein.

While the correlation between the two and its subsequent interpretation has been hotly debated, the general consensus in the field is that there is a clear correlation between the two (see Buccitelli & Selbach, Nature Reviews Genetics, 2020). Several reviews (Buccitelli and Selbach, 2020; Liu et al., 2016; Vogel and Marcotte,

2012) have suggested that two types of protein-mRNA correlations be distinguished. On one hand, the across-gene correlation can be estimated in the same sample across multiple proteins and the coding mRNAs. Alternatively, the within-gene correlation quantifies how the protein levels of a single gene correspond to the mRNA levels across multiple samples. Even though the within-gene correlations can vary greatly depending on the gene of interest and the conditions surveyed, recent studies conducted on mammalian tissues (Edfors et al., 2016; Moritz et al., 2019; Wang et al., 2019; Wilhelm et al., 2014) have consistently reported significant across-gene correlations, with an average Pearson correlation coefficient of 0.6 as summarized by a recent review (Buccitelli and Selbach, 2020). Our reanalysis of the proteogenomic data from human brains (Wang et al., 2019), found the mRNA and protein correlation for secretory pathway related genes to be around 0.7. The review also noted that true such correlations are expected to be higher due to the noise and bias inherent to the measurements (Buccitelli and Selbach, 2020). Here we found support score calculation relies on across-gene correlation since we calculate one support score for each individual using its condition-specific transcriptome, and no mRNA levels are used to approximate the protein abundance in a different cell or brain sample. Furthermore, variation in correlation between mRNA and protein for a given gene is expected to be balanced by other genes when comparing across samples, since the support score consists of component scores from many genes.

#### Choosing the number of eRWR iterations

The number of iterations was found empirically. Let $\pi$ be the steady-state (stationary) distribution for the eRWR. Suppose we reach steady state at time $t$, following the notations from the methods section the update rule becomes: $\mathbf{p}_{\mathbf{RWR}}(t) = (1-\alpha)\mathbf{W}_{\mathbf{adj}}\mathbf{p}_{\mathbf{RWR}}(\mathbf{t}) + \alpha\mathbf{p}(\mathbf{0})$. Rearranging terms, the stationary distribution can be derived as $\pi = \mathbf{p}_{\mathbf{RWR}}(t) = (\mathbf{I} - (1-\alpha)\mathbf{W}_{\mathbf{adj}})^{-1}\alpha\mathbf{p}(0)$. We varied the total iterations performed and plotted the convergence trajectories for the vector $\mathbf{p}_{\mathbf{RWR}}(t)$ on the inhibitory neuron-specific network. To facilitate the comparison, we divided each component $\mathbf{p}_{\mathbf{RWR}j}$ by its corresponding steady-state value $\pi_j$ so that a value of 1 indicates convergence. As is clear in Figure N3, 20 iterations of eRWR approximate the steady-state distribution rather well. In fact, at iteration $t$, the maximum total deviation from the steady can be upper bound by $\|\mathbf{p}_{\mathbf{RWR}}(t) - \pi\| \leq \sqrt{\frac{\max_a d(a)}{d(\mathrm{secP})}}\lambda_2^t$, where $d(a)$ indicates the degree of node $a$ and $\lambda_2$ the second-largest eigenvalue for the random walk transition matrix. Plugging in the values for our network, at $t = 20$ at least 98% convergence can be expected.

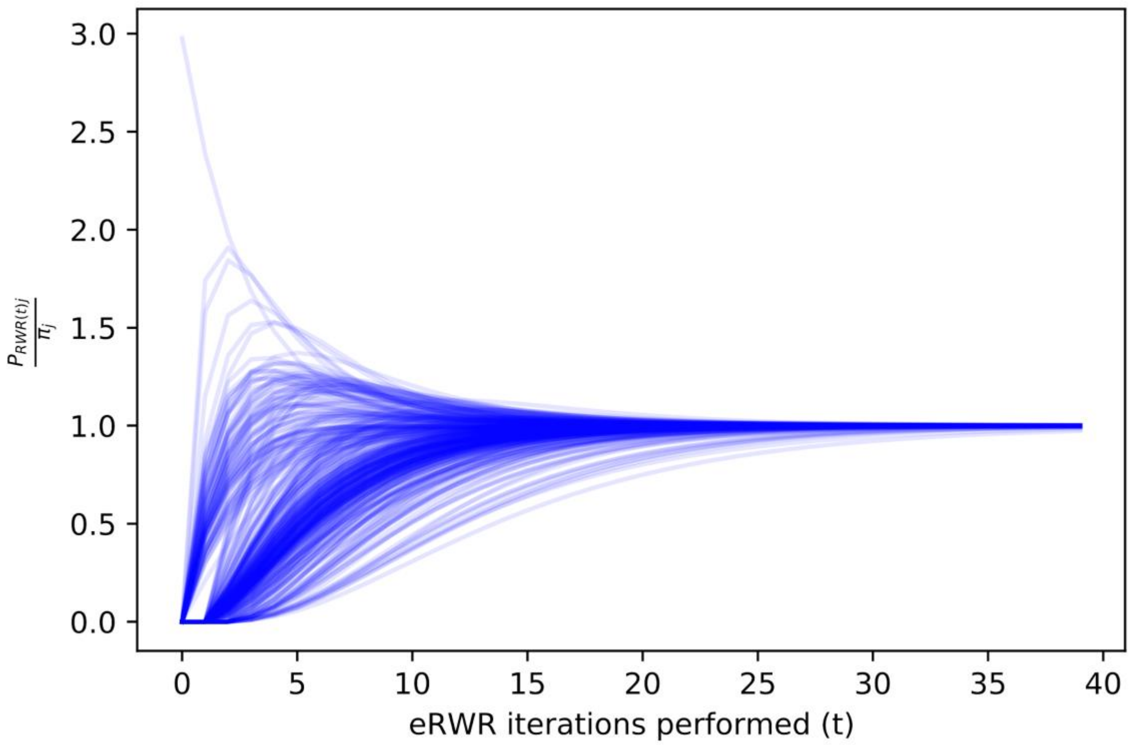


**Figure N3**
The support score component $\mathbf{p}_{\mathbf{RWR}_j}$ relative to its steady-state (stationary) distribution for each component (j) of the support network. Each line represents the relative convergence trajectory of one component at the $t^{th}$ eRWR iteration (x-axis).

#### Statistical analysis

##### Bayesian hierarchical model adaptations to account for sample covariates

The bulk RNA-seq dataset was previously corrected for known confounding factors including PMI, RACE, Batch, SEX, RIN and Exonic rate (Synapse ID syn20801188). For the ROSMAP snRNA-seq data, we expanded the model by incorporating the confounding factors including PMI, age and gender into the hierarchical model where available as suggested by the reviewer. We further imputed the age at death via a conditional censored model, since ages at death were censored (labeled 90+) for all individuals who died at the age of 90 or older. We additionally included the cell detection rate (CDR), which acts as a proxy for both technical (e.g., dropout, amplification efficiency) and biological factors (e.g., cell volume and extrinsic factors other than treatment of interest) that globally influence gene expression (Finak et al., 2015a). We calculated CDR for each single cell and added them to the Bayesian hierarchical model as a covariate. Figure S5 shows the distribution of all model parameters including the confounding factors.

##### Model performance: Tissue-specific expression of secMs and secPs

While the machinery support score improves the prediction of secP protein abundance from mRNA expression by >50% compared to the mRNA only model, we provide further evidence for the performance improvement. We summarized below the estimated the log marginal likelihoods $\log p(\mathrm{D}|M_i)$, along with out-of-sample predictive fit $\int_D \left( \log \int_\theta p(\widetilde{\mathrm{D}}_i|\theta) p_{\text{post}}(\theta)\, d\theta \right) f(\widetilde{\mathrm{D}}_i)\, d\widetilde{\mathrm{D}}$ (estimated by WAIC and scaled by a factor of -2 to convert from information criterion to predictive accuracy (Gelman et al., 2014)) for each model. The support score + mRNA model demonstrates significant improvement in both marginal likelihood given the dataset and the out-of-sample prediction accuracy, corresponding to a bayes factor of exp(477.62) and an improvement in expected predictive fit of exp(183.35).

| $M_i$ | ***log marginal likelihoods*** | ***WAIC/-2*** |
|---|---|---|
| *intercept + mRNA* | 19091.61 | -332437.2 |
| *intercept + mRNA + support score* | 19569.23 | -332253.85 |

##### Model performance: cross validation between multiple AD cohorts

The hierarchical model was performed on datasets where sequencing data with paired amyloid burden measurements are available. The only recent large-scale datasets meeting such criteria are the snRNA-RNAseq data from ROSMAP and bulk-RNAseq data from MSBB. We cross-validated the model trained from the MSBB dataset on the ROSMAP snRNA dataset. Caution must be taken when performing hierarchical model cross-validation between datasets with incompatible structures, such as the case for ROSMAP snRNA-seq and MSBB bulk-RNA-seq where data differ not only from the acquisition methods but also from their composition. To make the ROSMAP data compatible with the MSBB model, we randomly assigned the Broadmann Area to each single-cell data-point and median-summarized data points based on cell types. Since the Broadmann Area-associated effects are minor, the cross-validation accuracy is insensitive to this arbitrary assignment. We then predicted the amyloid levels for each subject--cell type combination based on the MSBB model posterior median. The cross-validation accuracies for neuronal cell types are the highest (R=0.46, 0.32; p = 0.0022, 0.037 for excitatory and inhibitory neurons respectively) among all cell types, in line with the results from the main text in which neurons showed the most significant support score-dependent amyloid deposition.

### Supplementary Figures

#### Figure S1

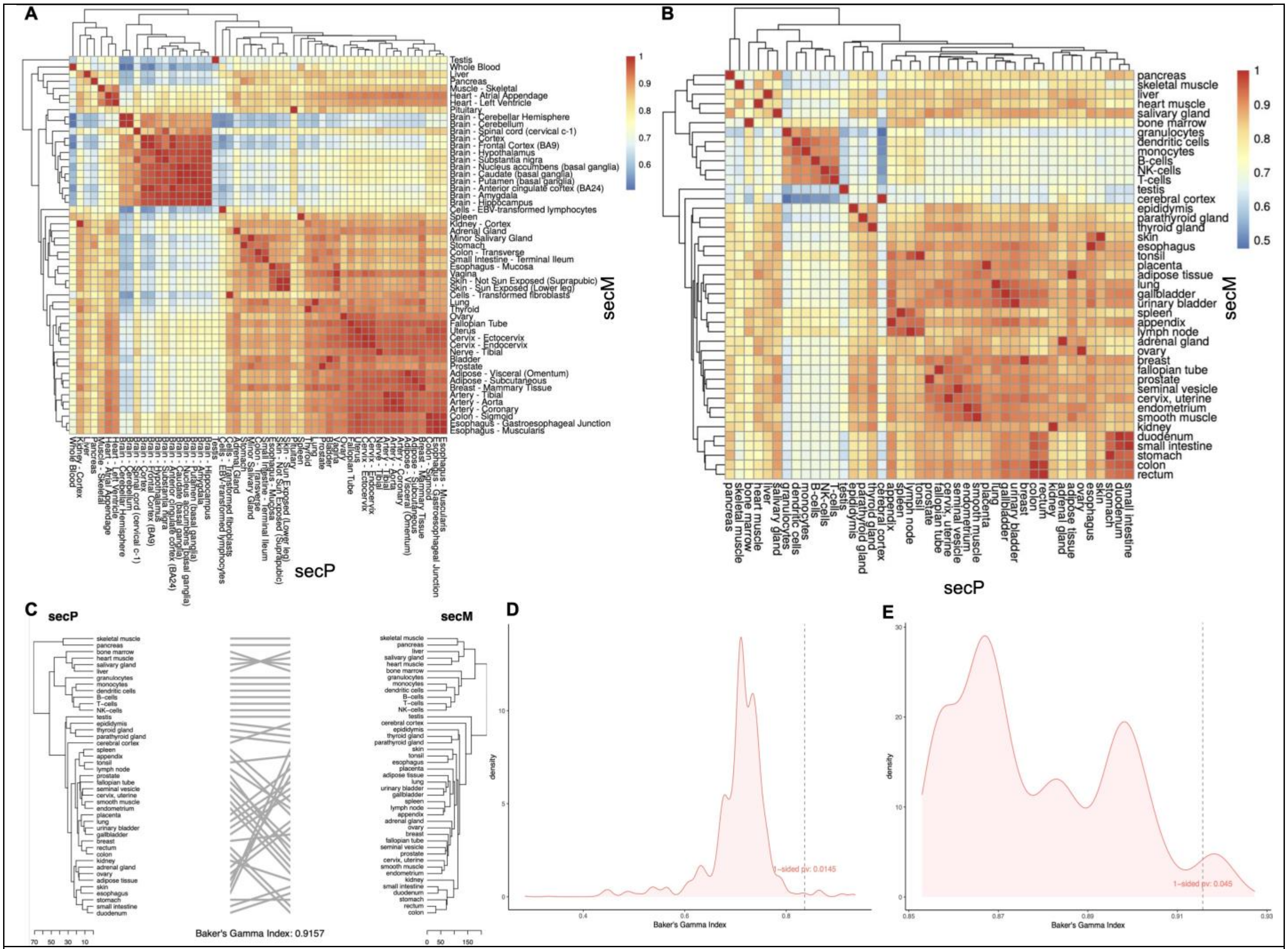


**The secMs and the secPs show coordinated expression profiles across different human tissues**

(A) GTEx (GTEx Consortium, 2015) Expression profile correlation matrix for clustered based on secPs (x-axis) and secMs (y-axis)

(B) Human Protein Atlas dataset (HPA) (Uhlén et al., 2015) Expression profile correlation matrix for clustered based on secPs (x-axis) and secMs (y-axis)

(C) Tanglegram (Galili, 2015) comparing the expression of the secretome and the secretory pathway expression across tissues from the HPA.

(D) The null permutation distribution for the Baker's Gamma index from the GTEx dataset obtained by randomly shuffling secMs with genesets of the same size. The permutation p-value is 0.0145.

(E) The null permutation distribution for the Baker's Gamma index from the Human Protein Atlas obtained by randomly shuffling secMs with genesets of the same size. The permutation p-value is 0.045.

### Figure S2

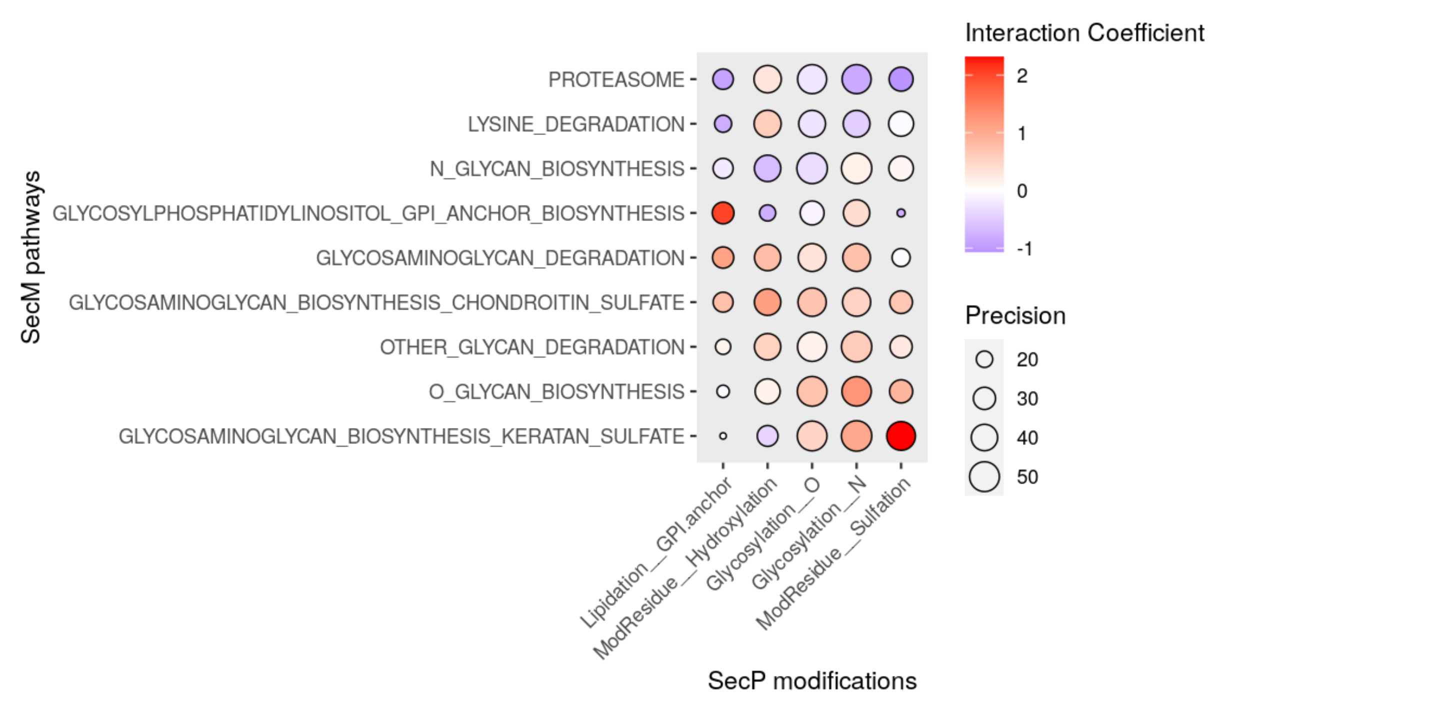


**PPIs between secPs and secMs recapitulate known protein post-translational modifications (PTMs).** The tendency for secM pathways to interact with secPs carrying certain PTMs. Positive interaction coefficients indicate more frequent interactions than expected by random chance. Several pairs of secM pathways and secP modifications with high interaction coefficients are between secPs harboring a given PTM and corresponding enzymes responsible for its synthesis, suggesting that many secPs have post-translational modifications that are added by specific proteins and that secPs typically have measured PPIs with the enzymes involved in their known PTMs. This provides evidence that the coordinated clustering across tissues for secPs and secMs could be driven by genes that are known to have PPIs.

The precision (inverse of variance) and the mean of the interaction coefficient are estimated with a linear interaction model detailed in our previous work on preferential protein interactions (Samoudi et al., 2020), and the precision of an interaction coefficient depends on the (estimated) strengths of the non-random association, the number of genes in secM each pathway and the ubiquity of the secP modification.

#### Figure S3

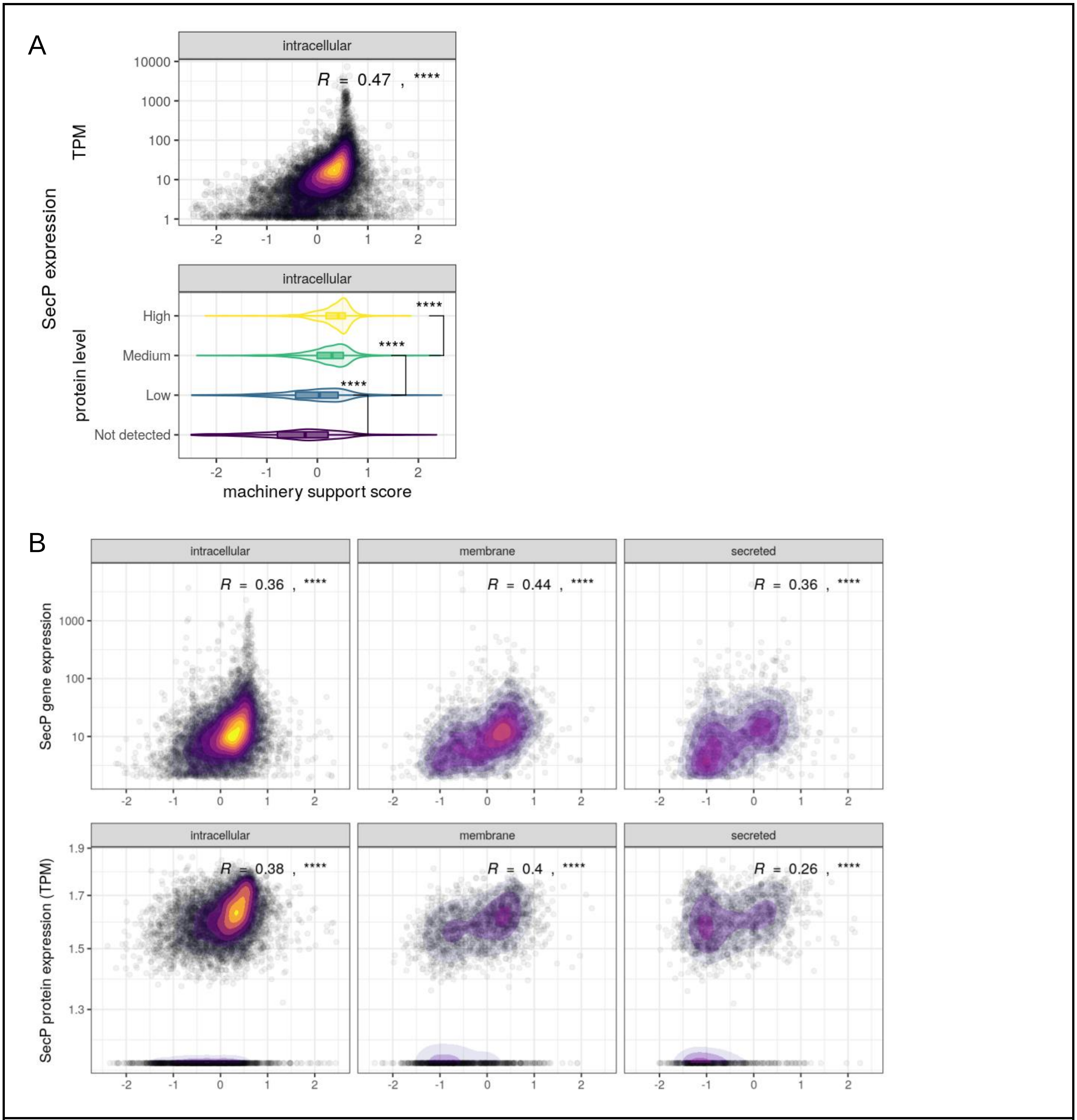


**Correlation between machinery support scores and transcript and protein abundance of secPs.**
(A) Relationship between the machinery support scores and the secP expression (top panel: transcript abundance, Spearman correlation coefficients; bottom panel: semi-quantitative protein abundance, t-test) of intracellular proteins from the Human Protein Atlas (Uhlén et al., 2015).
(B) Scatter plot comparing the machinery support scores and the secP expression (Spearman correlation coefficients. Top panel: transcript abundance; bottom panel: quantitative protein abundance) from the deep proteome and transcriptome abundance atlas (Wang et al., 2019).

#### Figure S4

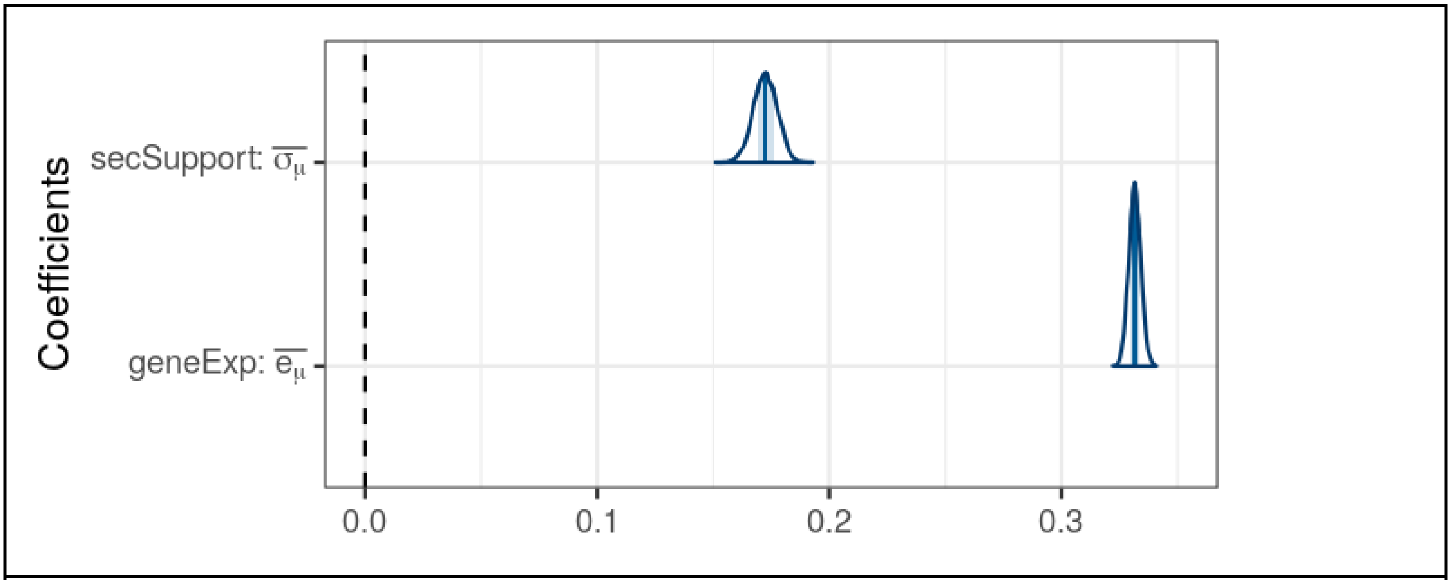


**Support scores predict secreted protein levels**
The posterior distribution of coefficients of protein support scores and gene expression in predicting protein abundance across tissues as estimated by the Bayesian hierarchical model (see methods).

#### Figure S5

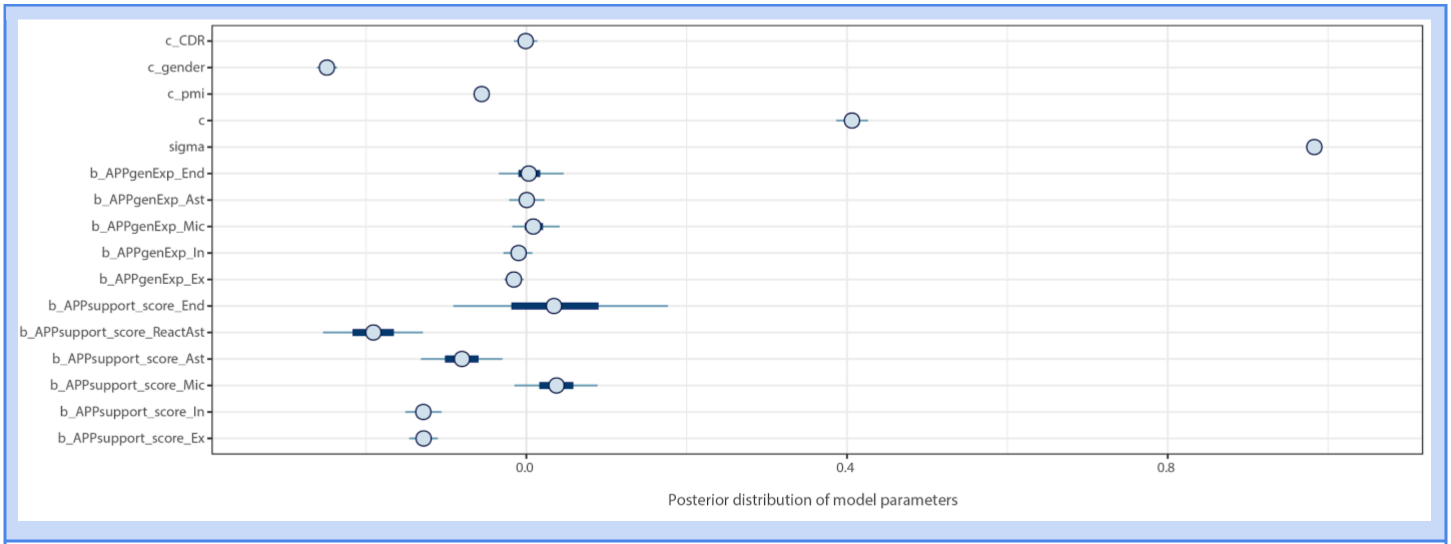


**The posterior distribution interval of all parameters in the cell-type specific model**
This figure shows all cell-type specific model parameters, of which Figure 3A plots a subset. CDR: cell detection rate (Finak et al., 2015b); PMI: post-mortem interval; End: endothelium; Ast: astrocytes; Mic: microglia; In: inhibitory neurons; Ex: excitatory neurons.

#### Figure S6

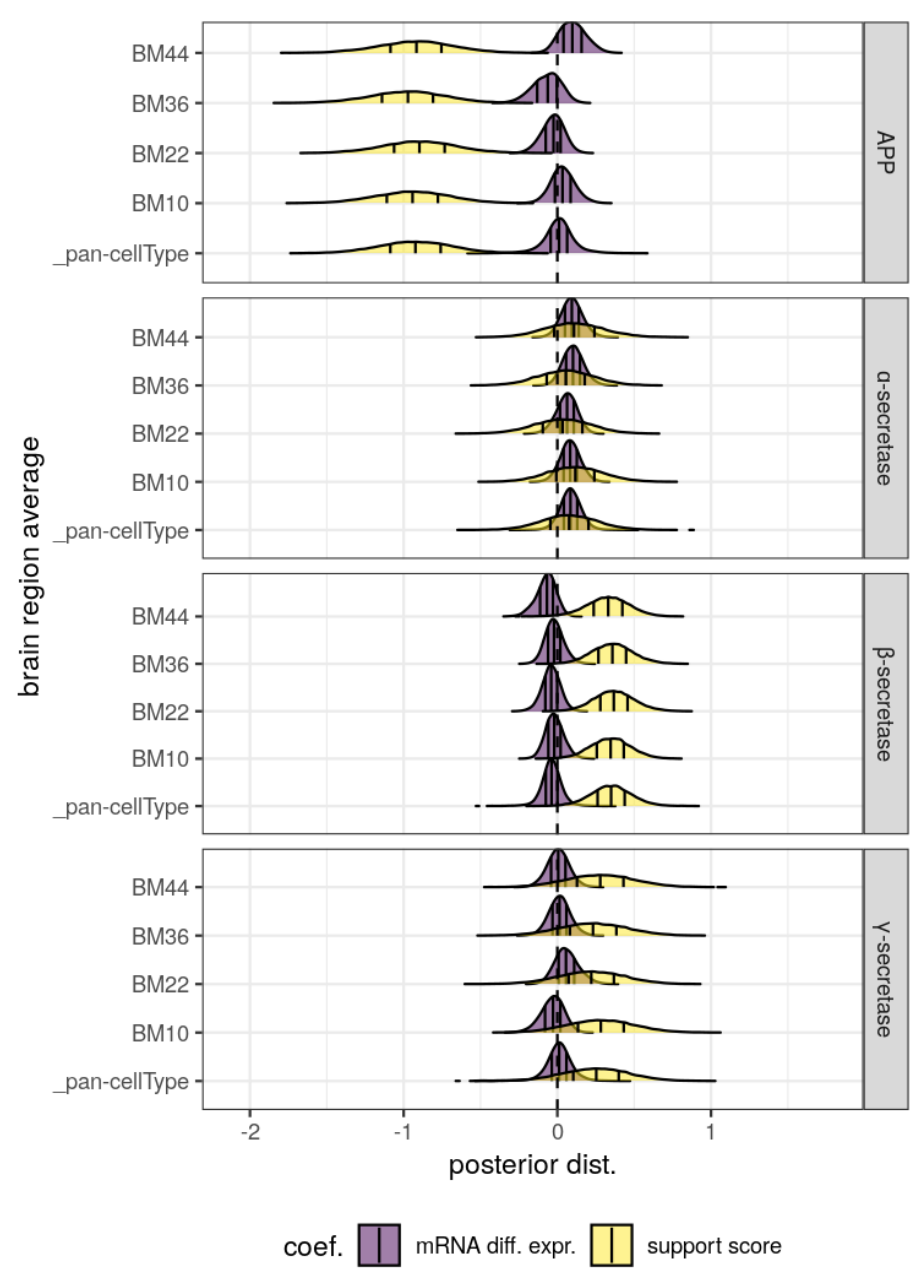


Posterior distribution of regression coefficients for mRNA differential expression and support scores for APP and the secretases by brain region. The brain region-average network is shown in Figure 3B.

Figure S7

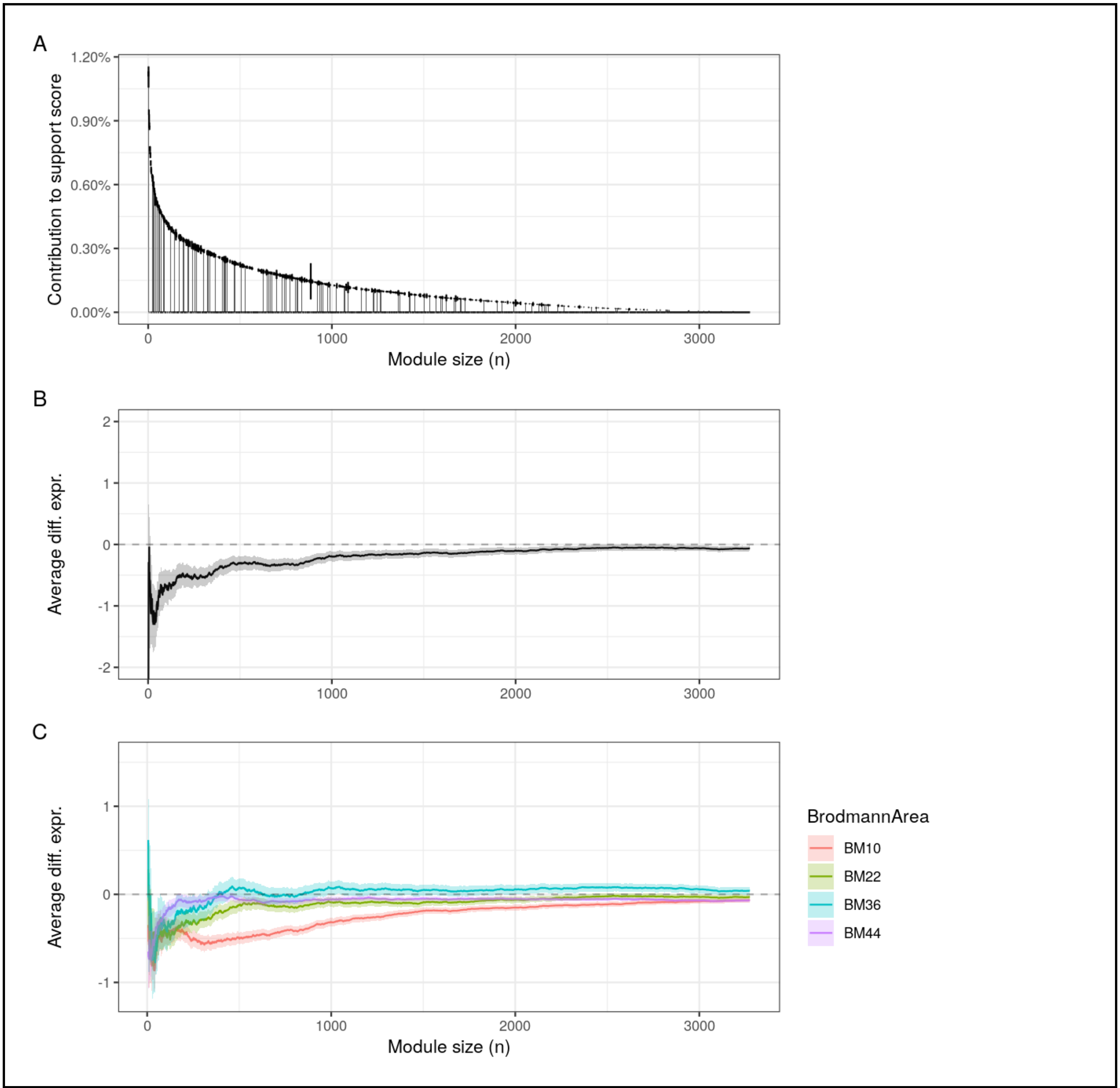


**(A)** Contribution to APP support score from each protein in the PPI network follows a power law. The component score for each protein in the support network (y-axis), rank-ordered by proximity to APP protein (x-axis, proteins most proximal to APP on the left), showed a pattern of exponential decay in which the support score is mainly determined by a few proteins.

**(B, C)** As an extension of Figures 4A and 4B, we show the overall (B) and brain region-specific (C) differential expression between AD and healthy subjects across all 3685 subnetworks.

#### Figure S8

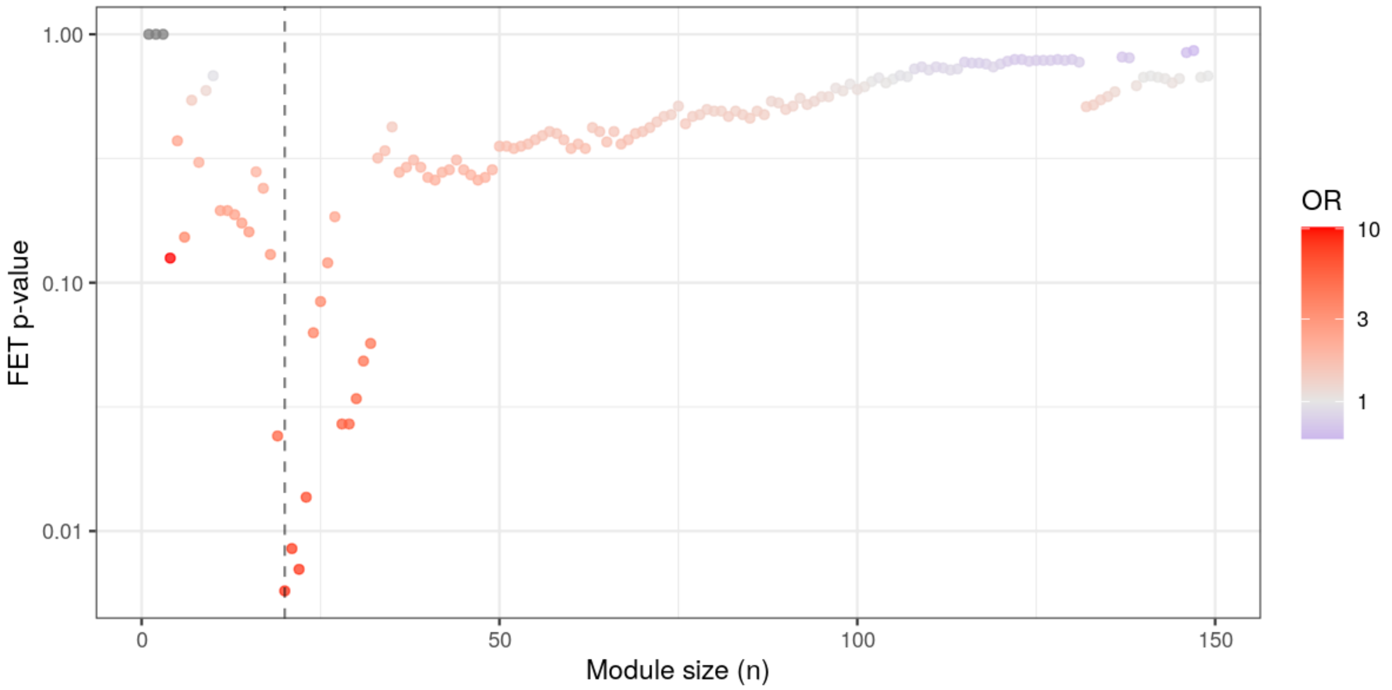


Corresponds to Figure 4B. Y-axis: FET p-value for the degree to which the subnetwork enriched for regulatory targets of known AD risk genes above the genome-wide significance threshold at each module size (x-axis).

138 # Figure S9

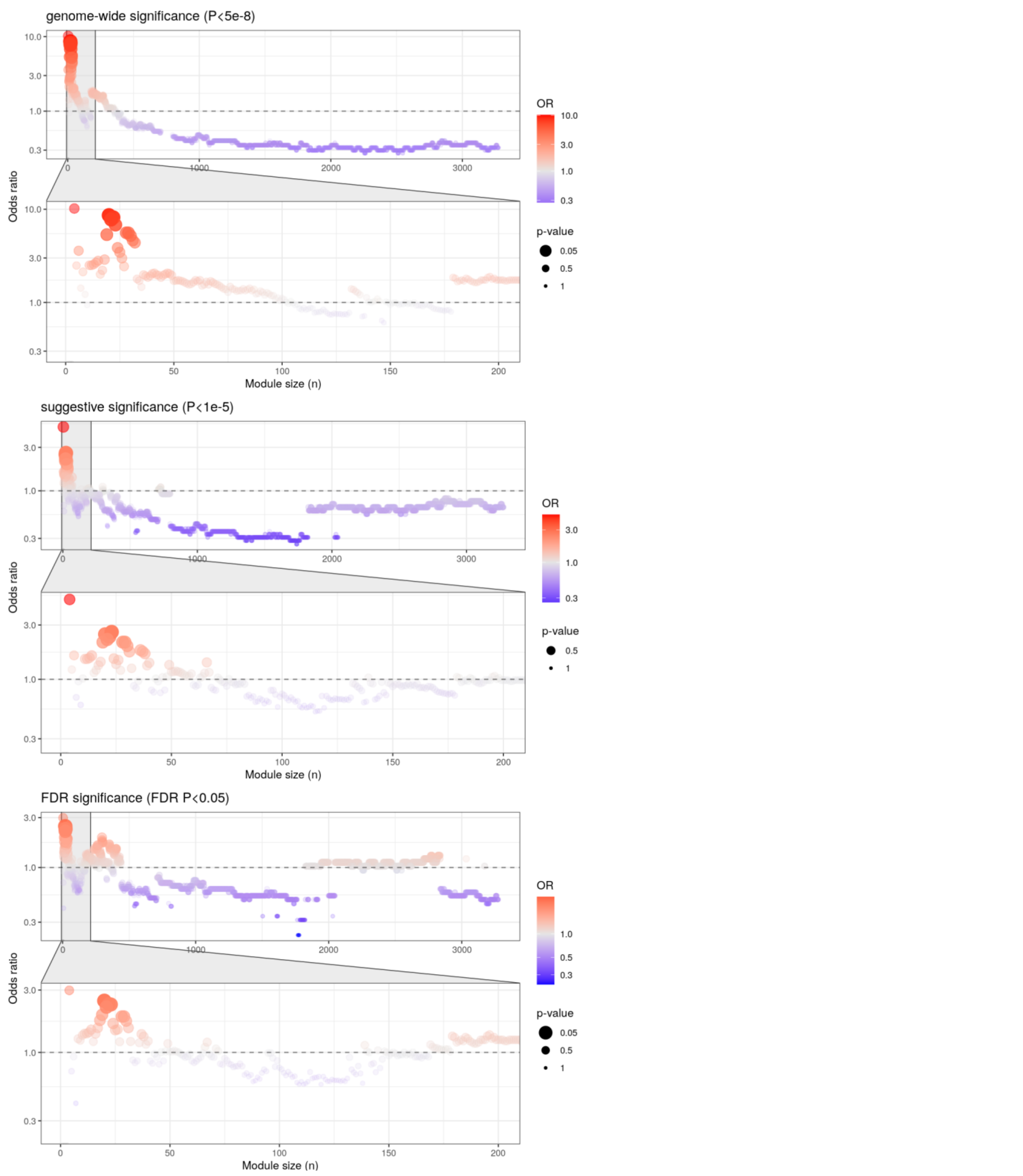


As an extension of Figure 4C, we show the degree to which the subnetworks enrich for regulatory targets of known AD risk genes above the genome-wide (top two panels, P<5e-8), suggestive (middle two panels, P<5e-7) and the FDR (bottom two panels, FDR P<.05) significance thresholds.

### Figure S10

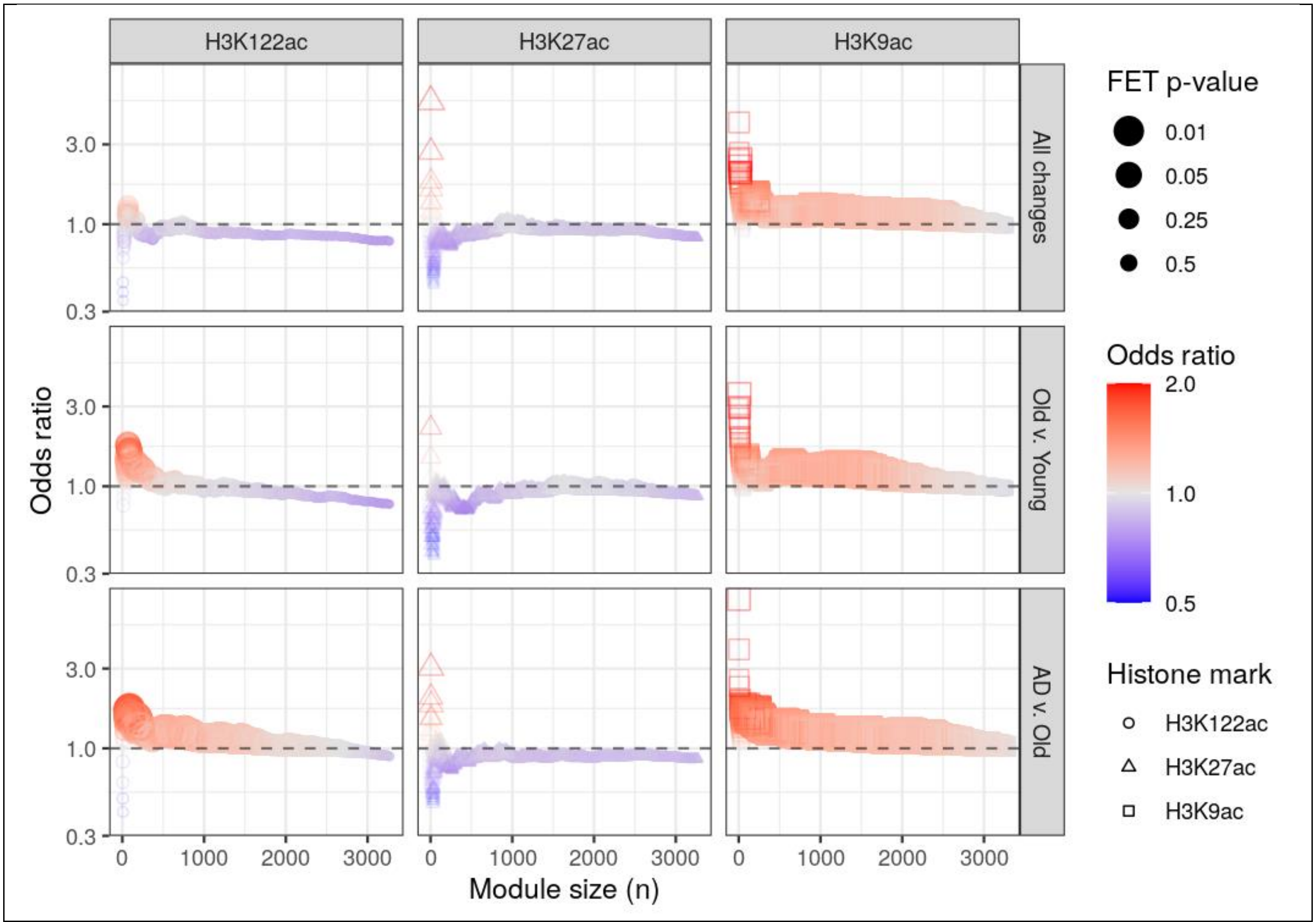


Following the notations from Figure 4, in each subplot the y-axis (odds ratio) shows the degree to which the core support network overlaps with genomic loci with differential histone modifications is indicated evaluated at modules of various sizes (x-axis), with a smaller n indicating a smaller subnetwork with only the proteins most proximal to APP. The subplots are arranged based on the histone marks profiled (columns), and the conditions compared (top row, age- or AD-related changes; middle row, age-specific changes; bottom row, AD-specific changes).

#### Figure S11

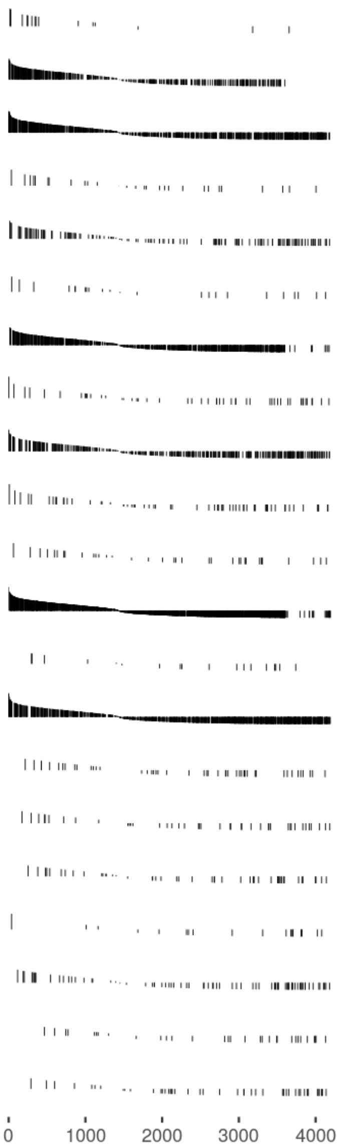


| Pathway | NES | pval | padj |
|---|---|---|---|
| Endosomes | 2.20 | 1.4e-04 | 1.1e-03 |
| Endoplasmic reticulum | 2.10 | 1.0e-04 | 1.1e-03 |
| Cytosol | 1.49 | 1.0e-04 | 1.1e-03 |
| Centriolar satellite | 1.42 | 9.9e-02 | 3.7e-01 |
| Nucleoli | 1.20 | 1.8e-01 | 4.2e-01 |
| Intermediate filaments | 1.04 | 4.3e-01 | 7.8e-01 |
| Golgi apparatus | 1.03 | 4.0e-01 | 7.8e-01 |
| Cell Junctions | 0.98 | 5.0e-01 | 7.8e-01 |
| Plasma membrane | 0.97 | 5.9e-01 | 7.8e-01 |
| Nuclear membrane | 0.90 | 6.2e-01 | 7.8e-01 |
| Microtubules | 0.87 | 6.3e-01 | 7.8e-01 |
| Vesicles | 0.86 | 9.7e-01 | 1.0e+00 |
| Cytokinetic bridge | 0.70 | 8.0e-01 | 9.6e-01 |
| Nucleoplasm | 0.66 | 1.0e+00 | 1.0e+00 |
| Centrosome | -0.78 | 8.5e-01 | 9.8e-01 |
| Nucleoli fibrillar center | -0.92 | 5.9e-01 | 7.8e-01 |
| Nuclear bodies | -1.27 | 1.1e-01 | 3.7e-01 |
| Focal adhesion sites | -1.37 | 1.2e-01 | 3.7e-01 |
| Mitochondria | -1.40 | 2.5e-02 | 1.1e-01 |
| Actin filaments | -1.61 | 2.2e-02 | 1.1e-01 |
| Nuclear speckles | -1.61 | 2.0e-02 | 1.1e-01 |

**Core support network is enriched for members of the amyloidogenic pathway.**
The component score for each member of the support network is compared with its stationary value in the network to quantify its relative proximity to APP, while controlling for network topology. With the secretory-resident proteins as the background, gene set enrichment analysis identified certain subcellular localizations to which proximal proteins are concentrated. This includes the ER, endosomes and the cytosol.

Figure S12

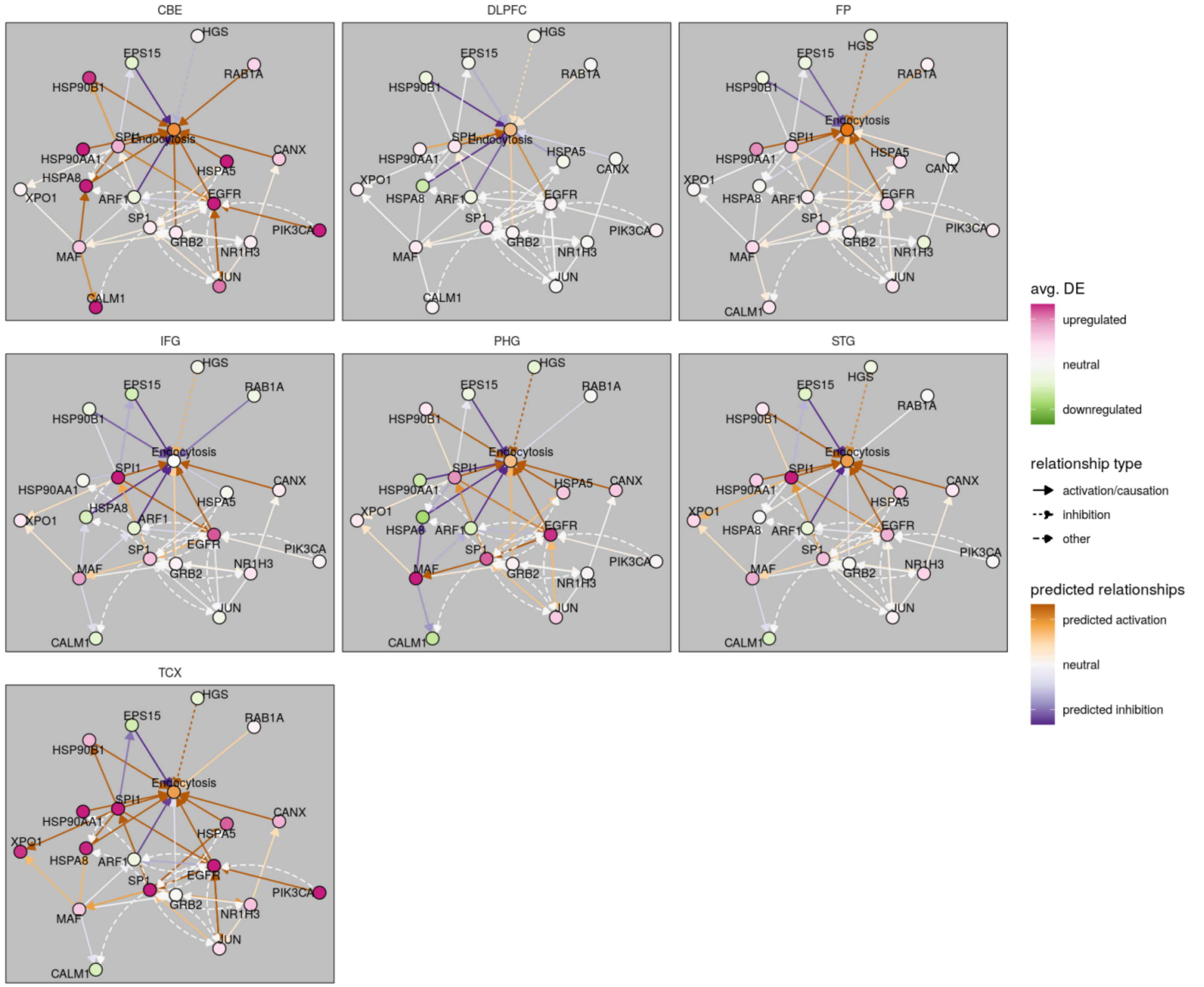


**The regulatory relationships surrounding the core support network with differential expression by brain regions-- Cerebellum(CBE), Dorsolateral prefrontal cortex (DLPFC), Frontal pole (FP, BM10), Inferior frontal gyrus (IFG, BM44), Parahippocampal gyrus (PHG, BM36), Superior temporal gyrus (STG, BM22) and Temporal cortex (TCX).**

TCX and CBE data were from the Mayo Study (Allen et al., 2016), DLPFC data were from ROSMAP study (Religious Order Study and Memory and Aging Project) (De Jager et al., 2018), and STG, FP, IFG, and PHG were from the Mount Sinai study (Wang et al., 2018). The predicted activation/ inhibition and network regulatory relationships were reported by the causal network analysis from Ingenuity Pathway Analysis (IPA) (Krämer et al., 2014). The brain region-average network is shown in Figure 5.

#### Figure S12

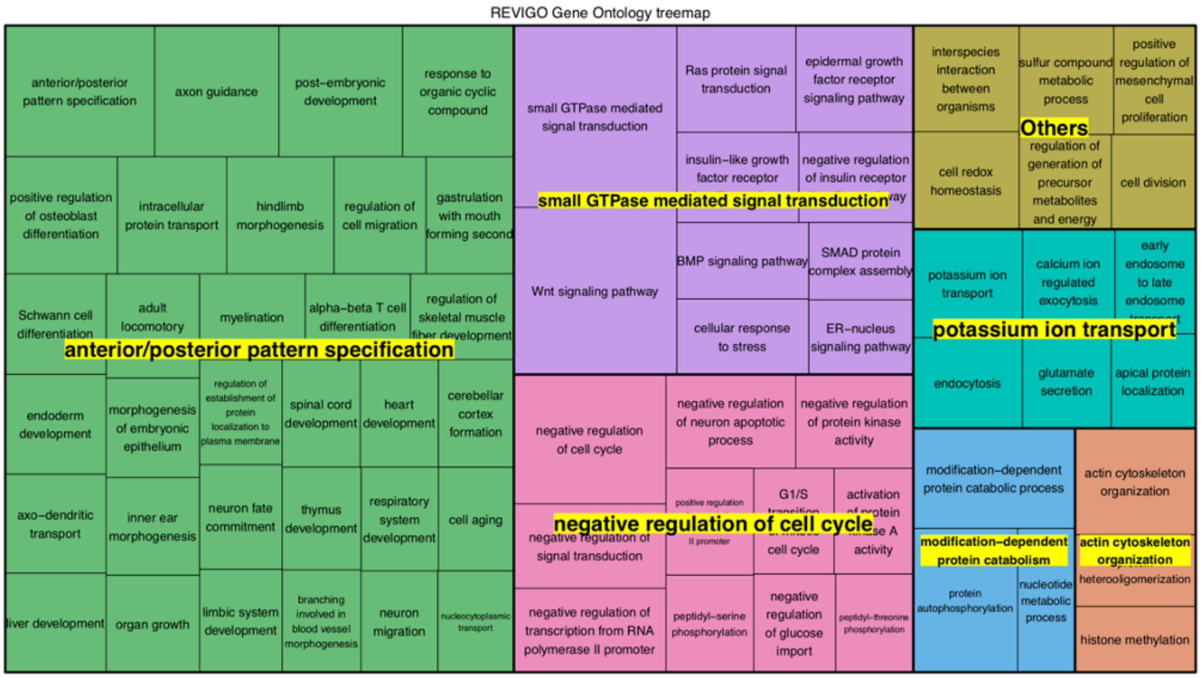


**GO functions associated with the conserved motif, as summarized by Revigo** (Supek et al., 2011)**.**
We found the conserved TF motif was significantly enriched in the promoter regions of known endocytic pathway genes (p-value=4.28e-4) and several other pathways relevant to AD pathogenesis, including anterior/posterior pattern specification(Damoiseaux et al., 2012), 2) small GTPase mediated signal transduction(Aguilar et al., 2017), 3) negative regulation of cell cycle(van Leeuwen and Hoozemans, 2015; Wu et al., 2016), 4) potassium ion transport(Cisternas et al., 2015), 5) modification-dependent protein catabolism(Ciavardelli et al., 2010; Gueli and Taibi, 2013), and 6) actin cytoskeleton organization(Hwang et al., 2016) (**Table S5**).

#### Supplementary Tables

##### Table S1 Proteome-wide secretory support scores for the HPA and the deep proteome

##### Table S2 Key amyloidogenic proteins Secretory support scores for the single-cell RNA-seq data from the ROSMAP project

The support scores received by key amyloidogenic proteins (APP, ADAM10, BACE1, and PSEN1) for each cell (indexed by *TAG*).

##### Table S3 Key amyloidogenic proteins Secretory support scores for the bulk-RNA-seq data (MSBB) from the ROSMAP project

The support scores received by key amyloidogenic proteins (APP, ADAM10, BACE1, and PSEN1) for each subject-brain region pair (indexed by *UID*).

##### Table S4 APP support component scores

The secretory component scores received by each protein (indicated by the *geneSymbol* column) in the APP support network. The component scores were averaged across subjects from the bulk-RNA seq (MSBB) data.

##### Table S5 Gene enhancers (GH) of NR1H3, MAF and SPI1 that related to global regulator -- SP1

##### Table S6 Go function enrichment for the TF motif


Aguilar, B.J., Zhu, Y., and Lu, Q. (2017). Rho GTPases as therapeutic targets in Alzheimer’s disease. Alzheimer’s Research & Therapy *9*.

Allen, M., Carrasquillo, M.M., Funk, C., Heavner, B.D., Zou, F., Younkin, C.S., Burgess, J.D., Chai, H.-S., Crook, J., Eddy, J.A., et al. (2016). Human whole genome genotype and transcriptome data for Alzheimer’s and other neurodegenerative diseases. Sci Data *3*, 160089.

Buccitelli, C., and Selbach, M. (2020). mRNAs, proteins and the emerging principles of gene expression control. Nat. Rev. Genet. *21*, 630–644.

Ciavardelli, D., Silvestri, E., Del Viscovo, A., Bomba, M., De Gregorio, D., Moreno, M., Di Ilio, C., Goglia, F., Canzoniero, L.M.T., and Sensi, S.L. (2010). Alterations of brain and cerebellar proteomes linked to Aβ and tau pathology in a female triple-transgenic murine model of Alzheimer's disease. Cell Death Dis. *1*, e90.

Cisternas, P., Lindsay, C.B., Salazar, P., Silva-Alvarez, C., Retamales, R.M., Serrano, F.G., Vio, C.P., and Inestrosa, N.C. (2015). The increased potassium intake improves cognitive performance and attenuates histopathological markers in a model of Alzheimer's disease. Biochim. Biophys. Acta *1852*, 2630–2644.

Damoiseaux, J.S., Prater, K.E., Miller, B.L., and Greicius, M.D. (2012). Functional connectivity tracks clinical deterioration in Alzheimer's disease. Neurobiol. Aging *33*, 828.e19–e828.e30.

De Jager, P.L., Ma, Y., McCabe, C., Xu, J., Vardarajan, B.N., Felsky, D., Klein, H.-U., White, C.C., Peters, M.A., Lodgson, B., et al. (2018). A multi-omic atlas of the human frontal cortex for aging and Alzheimer's disease research. Sci Data *5*, 180142.

Edfors, F., Danielsson, F., Hallström, B.M., Käll, L., Lundberg, E., Pontén, F., Forsström, B., and Uhlén, M. (2016). Gene-specific correlation of RNA and protein levels in human cells and tissues. Mol. Syst. Biol. *12*, 883.

Finak, G., McDavid, A., Yajima, M., Deng, J., Gersuk, V., Shalek, A.K., Slichter, C.K., Miller, H.W., McElrath, M.J., Prlic, M., et al. (2015a). MAST: a flexible statistical framework for assessing transcriptional changes and characterizing heterogeneity in single-cell RNA sequencing data. Genome Biol. *16*, 278.

Finak, G., McDavid, A., Yajima, M., Deng, J., Gersuk, V., Shalek, A.K., Slichter, C.K., Miller, H.W., McElrath, M.J., Prlic, M., et al. (2015b). MAST: a flexible statistical framework for assessing transcriptional changes and characterizing heterogeneity in single-cell RNA sequencing data. Genome Biol. *16*, 278.

Galili, T. (2015). dendextend: an R package for visualizing, adjusting and comparing trees of hierarchical clustering. Bioinformatics *31*, 3718–3720.

Gelman, A., Hwang, J., and Vehtari, A. (2014). Understanding predictive information criteria for Bayesian models. Stat. Comput. *24*, 997–1016.

Greenfield, J.P., Tsai, J., Gouras, G.K., Hai, B., Thinakaran, G., Checler, F., Sisodia, S.S., Greengard, P., and Xu, H. (1999). Endoplasmic reticulum and trans-Golgi network generate distinct populations of Alzheimer beta-amyloid peptides. Proc. Natl. Acad. Sci. U. S. A. *96*, 742–747.

GTEx Consortium (2015). Human genomics. The Genotype-Tissue Expression (GTEx) pilot analysis: multitissue gene regulation in humans. Science *348*, 648–660.

Gueli, M.C., and Taibi, G. (2013). Alzheimer's disease: amino acid levels and brain metabolic status. Neurol. Sci. *34*, 1575–1579.

Hartmann, T., Bieger, S.C., Brühl, B., Tienari, P.J., Ida, N., Allsop, D., Roberts, G.W., Masters, C.L., Dotti, C.G., Unsicker, K., et al. (1997). Distinct sites of intracellular production for Alzheimer's disease A beta40/42 amyloid peptides. Nat. Med. *3*, 1016–1020.

Huang, J.K., Carlin, D.E., Yu, M.K., Zhang, W., Kreisberg, J.F., Tamayo, P., and Ideker, T. (2018). Systematic Evaluation of Molecular Networks for Discovery of Disease Genes. Cell Syst *6*, 484–495.e5.

Huttlin, E.L., Ting, L., Bruckner, R.J., Gebreab, F., Gygi, M.P., Szpyt, J., Tam, S., Zarraga, G., Colby, G., Baltier, K., et al. (2015). The BioPlex Network: A Systematic Exploration of the Human Interactome. Cell *162*, 425–440.

Huttlin, E.L., Bruckner, R.J., Paulo, J.A., Cannon, J.R., Ting, L., Baltier, K., Colby, G., Gebreab, F., Gygi, M.P.,

Parzen, H., et al. (2017). Architecture of the human interactome defines protein communities and disease networks. Nature *545*, 505–509.

Hwang, J.Y., Shim, J.S., Song, M.-Y., Yim, S.-V., Lee, S.E., and Park, K.-S. (2016). Proteomic analysis reveals that the protective effects of ginsenoside Rb1 are associated with the actin cytoskeleton in β-amyloid-treated neuronal cells. J. Ginseng Res. *40*, 278–284.

Krämer, A., Green, J., Pollard, J., Jr, and Tugendreich, S. (2014). Causal analysis approaches in Ingenuity Pathway Analysis. Bioinformatics *30*, 523–530.

van Leeuwen, L.A.G., and Hoozemans, J.J.M. (2015). Physiological and pathophysiological functions of cell cycle proteins in post-mitotic neurons: implications for Alzheimer's disease. Acta Neuropathol. *129*, 511–525.

Liu, Y., Beyer, A., and Aebersold, R. (2016). On the Dependency of Cellular Protein Levels on mRNA Abundance. Cell *165*, 535–550.

Luck, K., Kim, D.-K., Lambourne, L., Spirohn, K., Begg, B.E., Bian, W., Brignall, R., Cafarelli, T., Campos-Laborie, F.J., Charloteaux, B., et al. (2020). A reference map of the human binary protein interactome. Nature *580*, 402–408.

Mathys, H., Davila-Velderrain, J., Peng, Z., Gao, F., Mohammadi, S., Young, J.Z., Menon, M., He, L., Abdurrob, F., Jiang, X., et al. (2019). Single-cell transcriptomic analysis of Alzheimer's disease. Nature *570*, 332–337.

Mohammadi, S., Davila-Velderrain, J., and Kellis, M. (2019). Reconstruction of Cell-type-Specific Interactomes at Single-Cell Resolution. Cell Syst *9*, 559–568.e4.

Moritz, C.P., Mühlhaus, T., Tenzer, S., Schulenborg, T., and Friauf, E. (2019). Poor transcript-protein correlation in the brain: negatively correlating gene products reveal neuronal polarity as a potential cause. J. Neurochem. *149*, 582–604.

Novick, P., Ferro, S., and Schekman, R. (1981). Order of events in the yeast secretory pathway. Cell *25*, 461–469.

Oughtred, R., Stark, C., Breitkreutz, B.-J., Rust, J., Boucher, L., Chang, C., Kolas, N., O'Donnell, L., Leung, G., McAdam, R., et al. (2019). The BioGRID interaction database: 2019 update. Nucleic Acids Res. *47*, D529–D541.

Reynaud, E.G., and Simpson, J.C. (2002). Navigating the secretory pathway: conference on exocytosis membrane structure and dynamics. EMBO Rep. *3*, 828–833.

Rolland, T., Taşan, M., Charloteaux, B., Pevzner, S.J., Zhong, Q., Sahni, N., Yi, S., Lemmens, I., Fontanillo, C., Mosca, R., et al. (2014). A proteome-scale map of the human interactome network. Cell *159*, 1212–1226.

Samoudi, M., Kuo, C.-C., Robinson, C.M., Shams-Ud-Doha, K., Schinn, S.-M., Kol, S., Weiss, L., Petersen Bjorn, S., Voldborg, B.G., Rosa Campos, A., et al. (2020). In situ detection of protein interactions for recombinant therapeutic enzymes. Biotechnol. Bioeng.

Supek, F., Bošnjak, M., Škunca, N., and Šmuc, T. (2011). REVIGO summarizes and visualizes long lists of gene ontology terms. PLoS One *6*, e21800.

Szklarczyk, D., Gable, A.L., Lyon, D., Junge, A., Wyder, S., Huerta-Cepas, J., Simonovic, M., Doncheva, N.T., Morris, J.H., Bork, P., et al. (2019). STRING v11: protein-protein association networks with increased coverage, supporting functional discovery in genome-wide experimental datasets. Nucleic Acids Res. *47*, D607–D613.

Uhlén, M., Fagerberg, L., Hallström, B.M., Lindskog, C., Oksvold, P., Mardinoglu, A., Sivertsson, Å., Kampf, C.,

Sjöstedt, E., Asplund, A., et al. (2015). Proteomics. Tissue-based map of the human proteome. Science *347*, 1260419.

Vogel, C., and Marcotte, E.M. (2012). Insights into the regulation of protein abundance from proteomic and transcriptomic analyses. Nat. Rev. Genet. *13*, 227–232.

Wang, D., Eraslan, B., Wieland, T., Hallström, B., Hopf, T., Zolg, D.P., Zecha, J., Asplund, A., Li, L.-H., Meng, C., et al. (2019). A deep proteome and transcriptome abundance atlas of 29 healthy human tissues. Mol. Syst. Biol. *15*, e8503.

Wang, M., Beckmann, N.D., Roussos, P., Wang, E., Zhou, X., Wang, Q., Ming, C., Neff, R., Ma, W., Fullard, J.F., et al. (2018). The Mount Sinai cohort of large-scale genomic, transcriptomic and proteomic data in Alzheimer's disease. Sci Data *5*, 180185.

Wilhelm, M., Schlegl, J., Hahne, H., Gholami, A.M., Lieberenz, M., Savitski, M.M., Ziegler, E., Butzmann, L., Gessulat, S., Marx, H., et al. (2014). Mass-spectrometry-based draft of the human proteome. Nature *509*, 582–587.

Wu, Y., Zhang, S., Xu, Q., Zou, H., Zhou, W., Cai, F., Li, T., and Song, W. (2016). Regulation of global gene expression and cell proliferation by APP. Sci. Rep. *6*, 22460.
